# Bifunctional Small Molecule Ligands of K-Ras Induce Its Association with Immunophilin Proteins

**DOI:** 10.1101/619551

**Authors:** Ziyang Zhang, Kevan M. Shokat

**Affiliations:** Dr. Z. Zhang, Prof. Dr. Kevan M. Shokat, Cellular and Molecular Pharmacology, Howard Hughes Medical Institute, University of California, San Francisco, 600 16th St. San Francisco, CA 94143

**Keywords:** Ras, cancer, drug design, immunophilin, dimerization

## Abstract

Here we report the design, synthesis and characterization of bifunctional chemical ligands that induce the association of Ras with ubiquitously expressed immunophilin proteins such as FKBP12 and cyclophilin A. We show this approach is applicable to two distinct Ras ligand scaffolds, and that both the identity of the immunophilin ligand and the linker chemistry affect compound efficacy in biochemical and cellular contexts. These ligands bind to Ras in an immunophilin-dependent fashion and mediate the formation of tripartite complexes of Ras, immunophilin and the ligand. The recruitment of cyclophilin A to GTP-bound Ras blocks its interaction with B-Raf in biochemical assays. Our study demonstrates the feasibility of ligand-induced association of Ras with intracellular proteins and suggests it as a promising therapeutic strategy for Ras-driven cancers.

## Introduction

Mutation of the Ras genes (K-, N-, and H-Ras) is the most common oncogenic lesion in cancer, occurring in more than 16% of all human cancers^[1]^. Absent deep accessible pockets, Ras proteins are often considered difficult to target directly by small molecule agents. Recent efforts have led to the identification of both peptidic^[2–6]^ and small-molecule^[7–14]^ direct binders of Ras. Targets of these binders include the nucleotide binding pocket, shallow hydrophobic patches, and dynamic pockets near the Switch-II region revealed only upon ligand binding (namely, the Switch-II pocket (S-IIP) and Switch-II groove (S-IIG)). While efficacious, the Switch-II pocket ligands face a few limitations: they depend on the presence of a nucleophilic Cys residue at position 12 and also require Ras to be in a GDP-bound (inactive) state^[11]^. Despite these limitations, several Switch-II pocket ligands have demonstrated potent cellular^[15]^, animal^[16]^ Ras inhibition and anti-tumor effects, and two compounds have entered clinical trials in K-Ras(G12C) mutant tumors. The Switch-II groove binding ligands, which target an unnatural M72C mutant are at an earlier stage of development exhibit the striking feature of recognizing both the GDP and GTP states and surprisingly block the association of PI3K but not Raf with drug-bound GTP-K-Ras (M72C)^[13]^. The limitations of these two classes of ligands bound to Switch II confound their use against several of the most frequent oncogenic mutations (e.g. G12D, G12V and Q61L), which compromise GTP hydrolysis and render Ras constitutively GTP-bound, and lack available cysteine residues. We wondered whether we could overcome these limitations by recruiting an endogenous protein to Ras with small molecule ligands. In so doing, we hoped to exploit the large surface area of the recruited protein to achieve high-affinity binding that might allow reversible ligand recognition of K-Ras and also introduce steric hinderance to block the interaction between Ras and its effectors.

Equipping an intracellular protein with a small molecule ligand to confer neo-functions is a concept employed by a number of natural and artificial systems. For example, rapamycin forms a complex with the PPIase FKBP12 and this complex binds the protein kinase mTOR to sterically block the access of its substrates^[17,18]^, while thalidomide induces the association of the E3 ligase Cereblon and the transcription factor Ikaros to promote the proteolytic degradation of the latter^[19,20]^. We envisioned that a bifunctional molecule could be constructed by chemically linking a Ras ligand and a high-affinity ligand of an abundant intracellular protein such as FKBP12 or cyclophilin A (CypA). FKBP12 has been reported to bind palmitoylated H-Ras and N-Ras and promote their depalmitoylation, thus we hoped that the two proteins might possess complementary surfaces naturally^[21]^. Here we report the design and synthesis of such bifunctional ligands and their successful induction of the association of Ras-FKBP12 or Ras-CypA. We show that both the structure of the linker and the nature of ligands chosen for FKBP12 or CypA impact the rate of ligand binding, and that a short piperazine linker endows the optimal efficiency among the compounds investigated.

## Results and Discussion

To design bifunctional molecules that recruit an intracellular protein to Ras, we first considered the identity of the intracellular protein. We chose two immunophilins, FKBP12 and CypA, as candidates because they are ubiquitously expressed chaperone proteins across all tissue types, and nature has repeatedly utilized them to target flat target surfaces upon binding of a small molecule agent (e.g. FK506^[22]^, rapamycin^[17,18]^, cyclosporin A^[22]^, and sanglifehrin A^[23]^). Several high-affinity ligands of FKBP12 and CypA have been reported (Fig. 1) and they each contain a solvent-exposed (when bound to FKBP12 or CypA) functional group handle that allows facile chemical derivatization. Next, we selected two chemical scaffolds as the Ras-binding component: Switch-II pocket ligands (S-IIP ligands, e.g. ARS1620 (**A**)) represent a family of covalent ligands that selectively binds the oncogenic Ras mutant G12C in the GDP-state^[11,15,16]^, whereas Switch-II groove ligands (S-IIG ligands, e.g. Compound 3 (**B**)) represent another class of covalent ligands that irreversibly reacts with a non-naturally occurring Ras mutant, M72C, regardless of the nucleotide binding state^[13]^. Both scaffolds occupy a pocket near the Switch-II region but have distinct binding modes. By analyzing co-crystal structures of these ligands bound to Ras, FKBP12 or CypA, we identified respective solvent-exposed sites for linker attachment that would likely preserve critical binding interactions. Finally, we devised a modular synthetic strategy that allowed us to independently vary three integral components (Ras ligand, FKBP12/CypA ligand and linker) to assemble the bifunctional ligands.

**Figure 1.**
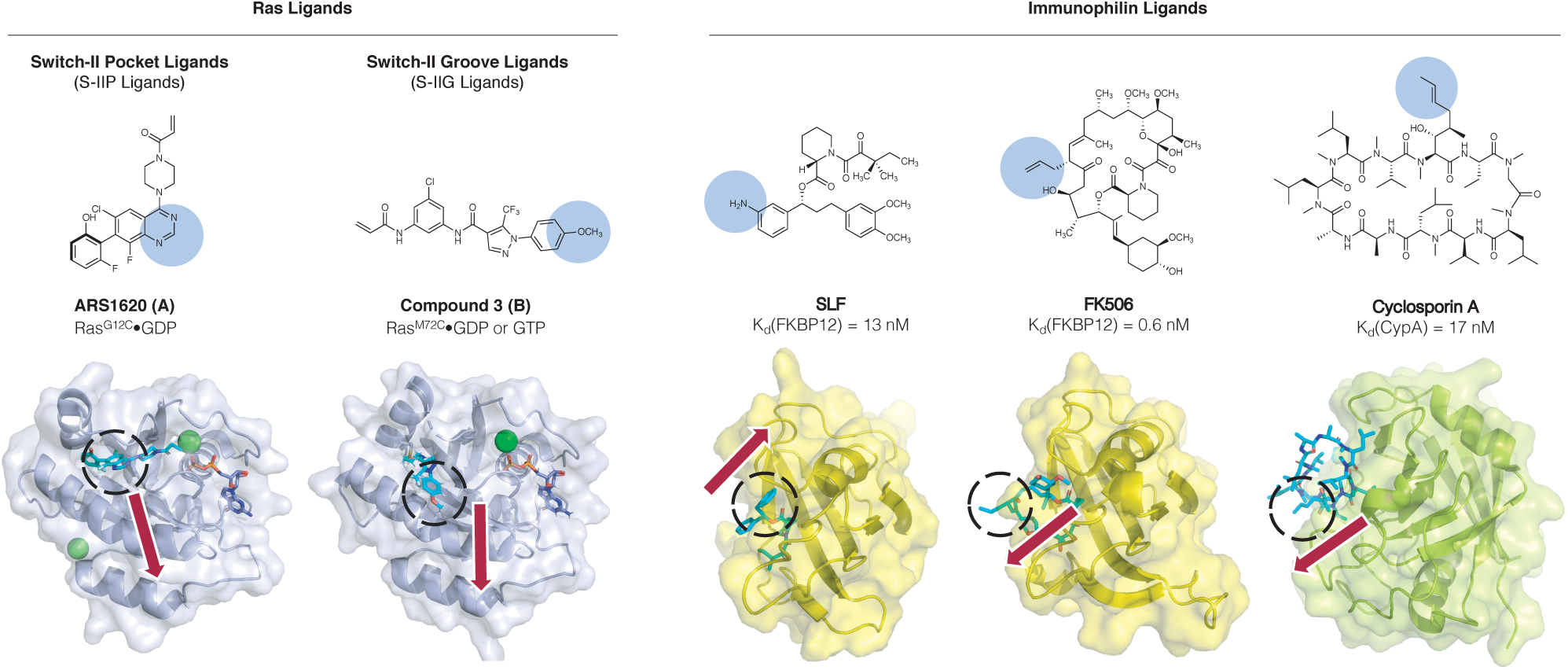
Small molecule ligands of Ras and immunophilins. Blue shades and dashed circles indicate the solvent-exposed sites of the ligands. Red arrows indicate the predicted trajectory of substituents attached at these sites. a) Chemical ligands of two distinct scaffolds that bind near the Switch II region of Ras. b) High-affinity ligands of FKBP12 (SLF and FK506) and of CypA (Cyclosporin A).

An initial series of compounds focused on the S-IIP scaffold were synthesized with various linker lengths and recruiter ligands including SLF, FK506 and cyclosporin A (Fig. 2a). All of these compounds maintained potent binding to FKBP12 or CypA (Kd < 200 nM, Supplementary Fig. 1). To evaluate the binding of these compounds to K-Ras, we measured the rate of their covalent labeling reaction with K-Ras G12C CysLight (1–169, C51S/C80L/C118S, GDP-bound) by intact protein mass spectrometry. Although the SLF-based compounds (**1**-**3**) failed to show substantial reactivity, both FK506-and cyclosporin-derived compounds (**4-6** and **7-10**, respectively) labeled K-Ras G12C CysLight at appreciable rates. Interestingly, including an excess (10 µM) of recombinant FKBP12 or CypA significantly accelerated the reactions of FK506-derived and cyclosporin A-derived compounds (Fig. 2b-d), respectively. By contrast, no change in reaction rates was observed with mismatched protein/compound combinations.

**Figure 2.**
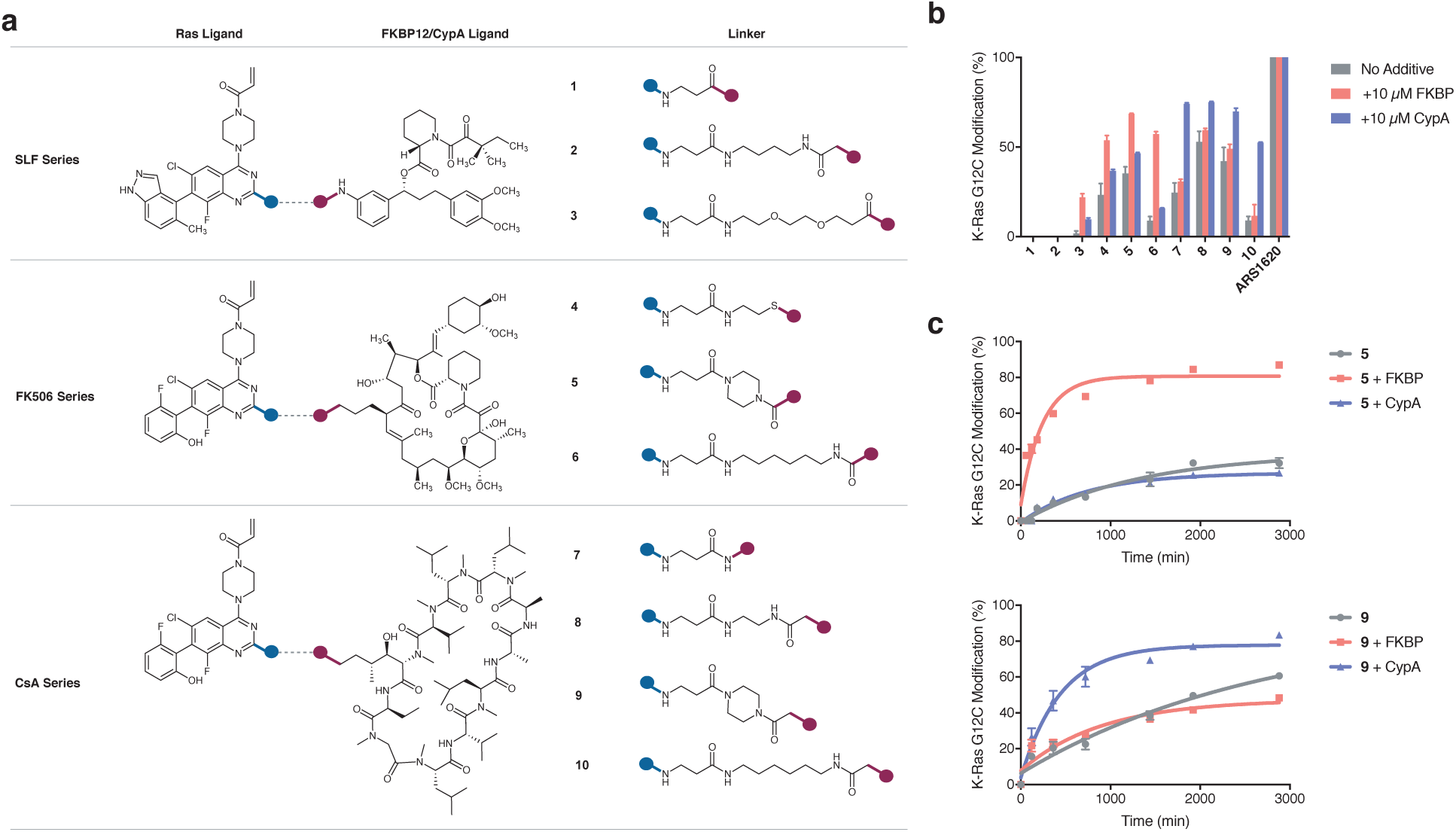
Bifunctional Ras(G12C) ligand created by linking a ligand of Ras and a ligand of FKBP12 or CypA. a) Modular design of bifunctional Ras ligands. b) Covalent modification of K-Ras(G12C) over 24 h at 23 °C. c) Immunophilin proteins accelerate the covalent labeling of K-Ras(G12C) by their cognate bifunctional molecules. All measurements were the average of three independent replicates.

In both FK506- and cyclosporin-derived series, compounds with a 1,6-diaminohexyl linker (**6** and **10**) demonstrated the slowest covalent adduct formation with K-Ras G12C CysLight. A plausible explanation is that long linear linkers impart more conformational flexibility and incur higher entropic cost upon compound binding. Though we have not established extensive structure-activity relationship, the trend we observed here is congruent with previous reports on bifunctional compounds designed for targeted protein degradation^[24]^.

To assess compound-induced association of Ras and immunophilins, we incubated a mixture of K-Ras G12C CysLight (4 µM), CypA (10 µM) and **8** (10 µM) at 23 °C for 24 h and confirmed >95% labeling of K-Ras by intact protein mass spectrometry. Size exclusion chromatography of the reaction mixture yielded a distinct major peak with a retention volume (Kav = 0.23) consistent with a ternary complex of the two proteins and the ligand at 1:1:1 stoichiometry (40 kDa), along with two smaller unresolved peaks corresponding to unreacted K-Ras and excess CypA (Fig. 3a). Differential scanning fluorimetry revealed two-stage thermal denaturation of the protein complex at 48.9 °C and 71.0 °C (Fig. 3b). By analysis of a protein sample heated at 60 °C for 10 min we confirmed that the first melting event corresponds to the denaturation of CypA (Fig. 3c), suggesting that the ligand, while greatly stabilizing K-Ras (Tm = 50.9 °C, Supplementary Fig. 2), did not increase the thermal stability of CypA (Tm = 49.1 °C, Supplementary Fig. 2).

**Figure 3.**
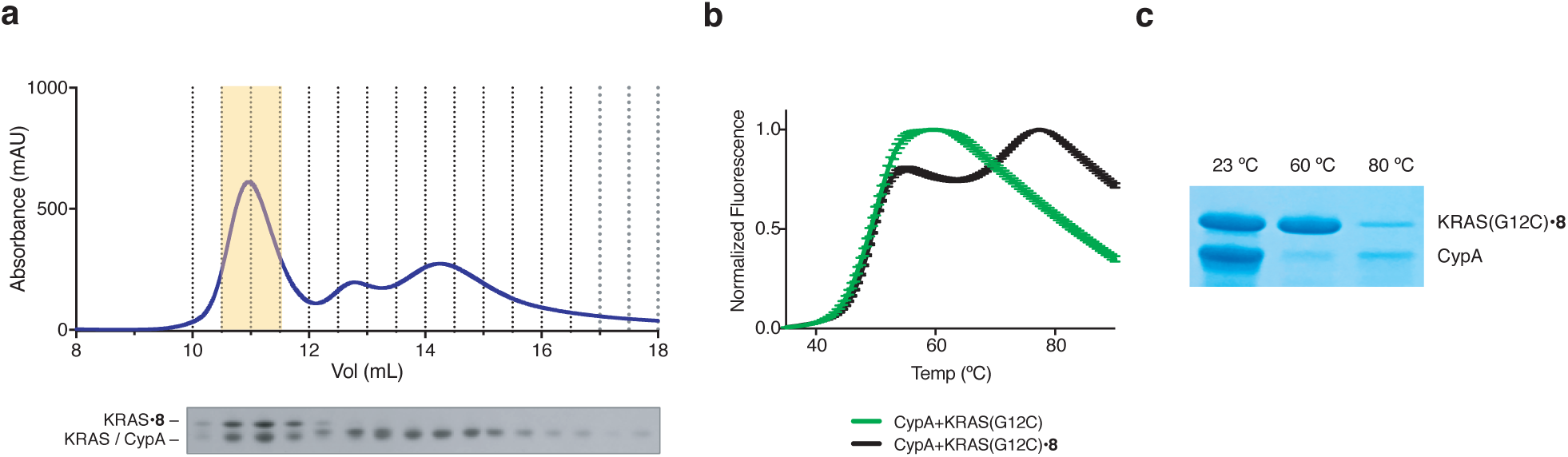
K-Ras(G12C) labeled by compound **8** forms a stable complex with CypA. a) Size exclusion chromatography of a reaction mixture consisting of K-Ras, **8** and cyclophilin A allows isolation of a 40-kDa protein complex. b) The K-Ras•**8**•CypA complex exhibits a two-step thermal denaturation profile. c) K-Ras•**8**•CypA complex was heated at 23 °C, 60 °C or 80 °C for 10 min and the soluble fraction was analyzed by SDS-PAGE.

We asked whether the recruitment of either CypA or FKBP12 would enable S-IIP ligands to interact with GTP-bound K-Ras and evaluated their ability to covalently react with K-Ras G12C CysLight (GppNHp-bound) in the presence of CypA or FKBP12. However, none of these molecules effected any detectable labeling of the GppNHp-bound protein even after 48 h of incubation at 23 °C, leading us to conclude that cysteine 12 remains inaccessible by this scaffold in the GppNHp-bound conformation of Ras^[16]^.

The cellular efficacy of the S-IIP bifunctional ligands were tested in NCI-H358, a cell line that harbors a heterozygous G12C mutation at the KRAS locus and has been shown to be sensitive to the G12C-selective covalent inhibitor ARS1620^[16]^. Partial covalent modification of Ras, revealed by the extra higher molecular weight bands in the immunoblot analysis with a pan-Ras antibody, was achieved by compounds **4-9** after 24-h incubation at 10 µM (Fig. 4a). The identity of these bands was confirmed by their selective pulldown using GST-FKBP12 or GST-CypA protein captured with glutathione agarose beads. Compounds **5** and **9**, both bearing a piperazine linker, led to the highest level of cellular target engagement in their respective series. Nevertheless, such a partial engagement appeared insufficient to impair the MAPK signaling pathways downstream of Ras, and the p-ERK levels remained unchanged, although we noticed a reduction of p-S6 level in **7**-and **9**-treated cells (Supplementary Fig. 3). Questioning whether the intracellular concentrations of CypA and FKBP12 were too low in this cell line to permit efficient recruitment by the compounds, we transiently transfected H358 cells with plasmids encoding HA-tagged FKBP12 or CypA proteins and treated the cells with **5** or **9** (Fig. 4b). Notwithstanding the expression of ectopic FKBP12 or CypA protein, we observed similar degrees of Ras modification and no change in p-ERK levels. We further considered that actively signaling Ras proteins are membrane-anchored and that both FKBP12 and CypA are cytosolic proteins, and hence investigated whether this discrepant sub-cellular distribution limited compound efficacy. With ectopic expression of plasma-membrane localized FKBP12 or CypA (each has a 11-mer N-terminal sequence derived from Lyn to direct plasma-membrane localization^[25]^), we observed compound-dependent attenuation of p-ERK signal (Fig. 4b), despite that the level of Ras modification was comparable to that in mock-transfected cells. While further investigation is merited to elucidate the exact Ras sub-population responsible for this change, we believe that these membrane-bound immunophilins directly improved compound efficacy via their localized expression. Together, these results confirmed that a high level of target engagement of membrane-bound K-Ras G12C is essential to impede its signaling output through the MAPK pathway, and suggest that further optimization on these compounds is necessary to improve their cellular activity.

**Figure 4.**
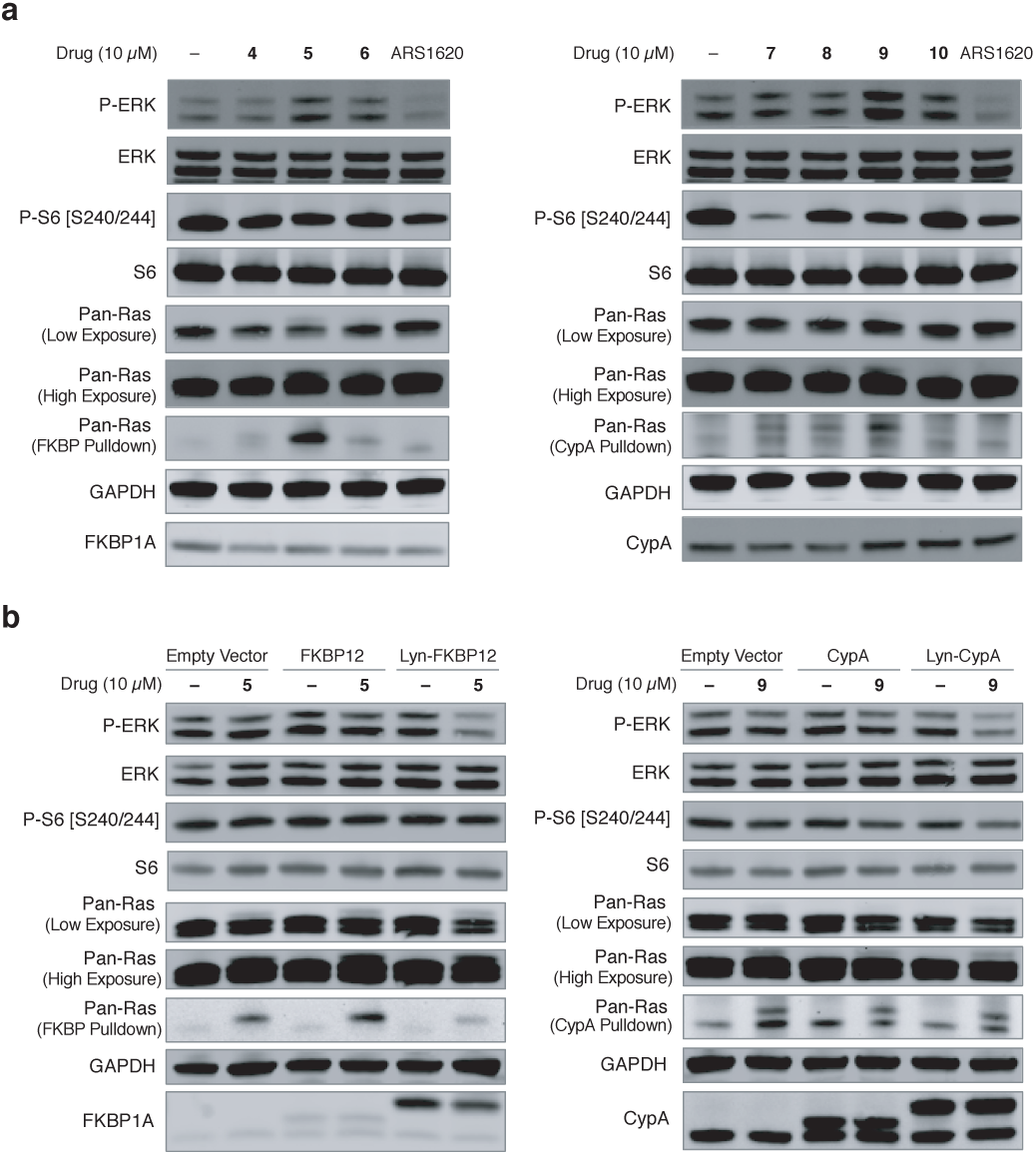
Bifunctional K-Ras(G12C) ligands have limited efficacy against H358, a cell line harboring a G12C mutation in one KRAS allele. a) H358 cells were treated with DMSO or compounds for 24 h and analyzed by immunoblotting. b) HA-tagged FKBP12, Lyn-FKBP12, CypA, or Lyn-CypA was ectopically expressed in H358 cells by transient transfection, then the cells were treated with DMSO or indicated compounds for 24 h and analyzed by immunoblotting. The presented data is representative of three independent replicates.

In order to access the GTP-bound state of Ras, we explored a different scaffold (S-IIG ligands) that has been demonstrated to bind both the GDP- and GTP-bound states of Ras harboring a non-natural mutation M72C (Fig. 1a). When covalently bound to Ras, compounds derived from this scaffold inhibit the interaction of Ras with its nucleotide exchange factor Sos and its downstream signal transducer PI3K, but the structural changes induced by compound binding were insufficient to block the binding of Raf, the Ras effector that transduces signaling along the MAPK pathway. To enable recruitment of FKBP12 or CypA to GTP-bound Ras, we synthesized FK506-based compound **11** and cyclosporin A-based compound **12** using a similar synthetic route (Fig. 5a). While the M72C mutation was initially introduced to guide site-directed ligand discovery, it also served as a useful tool to evaluate compound binding. Analogous to S-IIP bifunctional ligands, **11** and **12** displayed FKBP- and CypA-accelerated labeling of K-Ras M72C CysLight (GDP-bound or GppNHp-bound), and the cyclosporin-derived compound **12** appeared superior to the FK506-derived **11** in its rate of covalent labeling (Fig. 5b). Ternary complexes consisting of K-Ras M72C CysLight (GDP-or GppNHp-bound), CypA and **12** were purified (Supplementary Fig. 4) and used to study whether the induced associated of CypA and Ras is able to block the binding of Ras effectors such as Sos and Raf.

**Figure 5.**
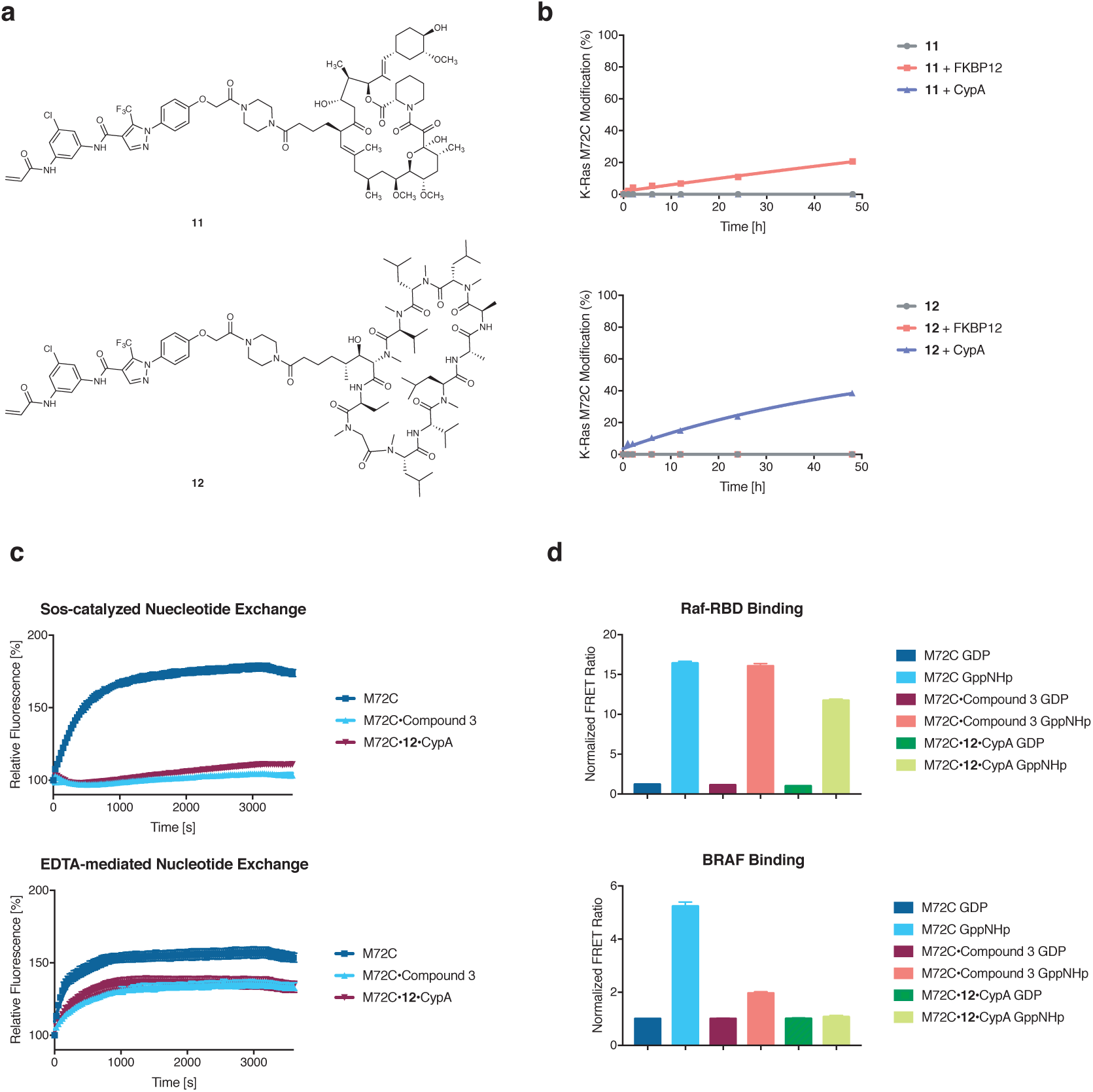
Bifunctional Ras(M72C) ligands and their biochemical characterization. a) Chemical structures of two bifunctional Ras(M72C) ligands. b) Compounds **11** and **12** covalently react with K-Ras(M72C) in an FKBP12/CypA-dependent fashion. c) The K-Ras•**12**•CypA complex is resistant to Sos-catalyzed but not EDTA-mediated nucleotide exchange reaction. d) The K-Ras•**12**•CypA complex retains the ability to bind Raf-RBD but not full length BRAF protein. All measurements are the average of three independent replicates.

Similar to the adduct formed between K-Ras M72C and S-IIG ligand **B**^[13]^, the K-Ras M72C•**12**•CypA complex was insensitive to Sos-catalyzed nucleotide exchange but susceptible to the EDTA-mediated reaction (Fig. 5c). To investigate the effect of the associated CypA on Raf binding, we employed a solution-based time-resolved fluorescence resonance energy transfer (TR-FRET) assay that allows the quantitation of the Ras•Raf complex formation. We compared K-Ras M72C, K-Ras M72C•**B** and K-Ras M72C•**12**•CypA in both GDP-and GppNHp-bound forms in terms of their ability to form a complex with the GST-tagged Ras-binding domain of Raf1 (Raf1-RBD). While none of the GDP-bound proteins had detectable interaction with Raf1-RBD, all the three GppNHp-bound forms led to a dramatic increase in the TR-FRET signal (Fig 5d). Studying the published crystal structures of the H-Ras•Raf1-RBD complex and of H-Ras bound by a closely related ligand 2C07^[13]^ revealed the possibility that the recruited CypA protein may adopt an orientation to position itself away from Switch I, where the key interactions between Ras and Raf1-RBD reside. In such a scenario, Ras may simultaneously bind Raf1-RBD and CypA without inflicting a steric clash of the two proteins. Considering the much larger sizes of native Raf proteins than the truncated Raf-RBD, we wondered if full length B-Raf protein would be a more appropriate means to study Ras•Raf interaction^[26,27]^. When the same assay was performed using recombinant GST-B-Raf protein, we observed a compound-dependent effect on Raf binding: compound **B**-labeled K-Ras showed weakened association with full length B-Raf, whereas the K-Ras•**12**•CypA complex completely lost this interaction (Fig. 5d). This decrease in affinity between the ternary complex and B-Raf supports our hypothesis that CypA can act as a steric blocker of Ras-effector communication.

## Conclusion

Recent discoveries of direct small molecule ligands of Ras have transformed the therapeutic prospects of Ras-driven cancer ^[3,5,7–10,14]^. Switch-II pocket and Switch-II groove binders reveal new opportunities to inhibit Ras, and the former have demonstrated efficacy in xenograft models for K-RAS(G12C)-driven cancer and are currently in clinical trials^[16]^. Meanwhile, three key challenges remain: 1) the ability to access the GTP-bound state of Ras is essential to inhibiting many oncogenic mutants of Ras that are constitutively GTP-bound; 2) ligands must be able to block the interaction between Ras•GTP and multiple effector proteins including Raf and PI3K to have therapeutic effect; and 3) the ability to discover non-cysteine dependent ligands to target K-Ras. Neither current S-IIP nor S-IIG ligands meet all of these criteria. While it is conceivable to overcome these limitations with extensive structure based medicinal chemistry optimization of current ligands, we sought to draw inspiration from the unique mode of action of drugs that induce the formation of ternary protein-drug-protein complexes^[28,29]^ and to develop small molecules that recruit intracellular proteins to block Ras-effector interactions.

Our study showed that small molecule-induced association of K-Ras and immunophilins is possible even though these proteins do not have measurable intrinsic affinity for each other. The complete blockade of Ras-Raf and Ras-Sos interaction by the combination of compound **12** and CypA illustrates the power of steric hinderance at insulating Ras from its effectors and affirms the potential of compound-induced recruitment of intracellular proteins as a strategy to inhibit protein-protein interactions. We acknowledge that the compounds discussed in the present work do not possess the necessary efficacy for detailed cellular studies, and that the S-IIG ligands are yet dependent on their covalent reactivity with a non-natural cysteine (M72C). However, we believe our characterization of the structure activity relationships across two Ras ligands and three immunophilin ligands as well as multiple linkers provides a starting point for further optimization. Importantly, our studies using whole protein mass spectrometry to detect K-Ras (G12C) binding in vitro as well as cellular engagement assessment by immunophilin pull-down followed by anti-Ras immunoblot showed good correlation between biochemical and cellular target engagement, providing a path for further structural optimization. Further chemical optimization will be necessary to improve the affinity of current ligand scaffolds for Ras. Perhaps new ligands for Ras directed at other surface pockets and grooves when they are reported will be attractive directions. Nevertheless, these compounds offered key evidence for the feasibility of our approach, and we are confident that with additional efforts, obstructing the signaling output of hyperactive oncogenic Ras mutants with intracellular proteins will become a promising therapeutic strategy.

## Acknowledgements

The authors would like to thank Dr. Roger Briesewitz for his invaluable advice and encouragement throughout all stages of the project, Dr. Ming Sun and Dr. Adam Frost for their help with structural work. ZZ is a Damon Runyon HHMI Fellow supported by the Damon Runyon Cancer Research Foundation (DRG-2281-17).

## References

[1] I. A. Prior, P. D. Lewis, C. Mattos, Cancer Res. 2012, 72, 2457–2467.

[2] S. Nickerson, S. T. Joy, P. S. Arora, D. Bar-Sagi, Nat. Chem. Biol. 2011, 7, 585–587.

[3] T. B. Trinh, P. Upadhyaya, Z. Qian, D. Pei, ACS Comb. Sci. 2016, 18, 75–85.

[4] R. Spencer-Smith, A. Koide, Y. Zhou, R. R. Eguchi, F. Sha, P. Gajwani, D. Santana, A. Gupta, M. Jacobs, E. Herrero-Garcia, et al., Nat. Chem. Biol. 2017, 13, 62–68.

[5] S. Sogabe, Y. Kamada, M. Miwa, A. Niida, T. Sameshima, M. Kamaura, K. Yonemori, S. Sasaki, J. Sakamoto, K. Sakamoto, ACS Med. Chem. Lett. 2017, 8, 732–736.

[6] J. H. McGee, S. Y. Shim, S. J. Lee, P. K. Swanson, S. Y. Jiang, M. A. Durney, G. L. Verdine, J. Biol. Chem. 2018, 293, 3265–3280.

[7] A. G. Taveras, S. W. Remiszewski, R. J. Doll, D. Cesarz, E. C. Huang, P. Kirschmeier, B. N. Pramanik, M. E. Snow, Y. S. Wang, J. D. Del Rosario, et al., Bioorganic Med. Chem. 1997, 5, 125–133.

[8] T. Maurer, L. S. Garrenton, A. Oh, K. Pitts, D. J. Anderson, N. J. Skelton, B. P. Fauber, B. Pan, S. Malek, D. Stokoe, et al., Proc. Natl. Acad. Sci. U. S. A. 2012, 109, 5299–304.

[9] Q. Sun, J. P. Burke, J. Phan, M. C. Burns, E. T. Olejniczak, A. G. Waterson, T. Lee, O. W. Rossanese, S. W. Fesik, Angew. Chem. Int. Ed. 2012, 51, 6140–6143.

[10] F. Shima, Y. Yoshikawa, M. Ye, M. Araki, S. Matsumoto, J. Liao, L. Hu, T. Sugimoto, Y. Ijiri, A. Takeda, et al., Proc. Natl. Acad. Sci. 2013, 110, 8182–8187.

[11] J. M. Ostrem, U. Peters, M. L. Sos, J. A. Wells, K. M. Shokat, Nature 2013, 503, 548–51.

[12] S. M. Lim, K. D. Westover, S. B. Ficarro, R. A. Harrison, H. G. Choi, M. E. Pacold, M. Carrasco, J. Hunter, N. D. Kim, T. Xie, et al., Angew. Chem. Int. Ed. 2014, 53, 199–204.

[13] D. R. Gentile, M. K. Rathinaswamy, M. L. Jenkins, S. M. Moss, B. D. Siempelkamp, A. R. Renslo, J. E. Burke, K. M. Shokat, Cell Chem. Biol. 2017, 24, 1455–1466.e14.

[14] M. E. Welsch, A. Kaplan, J. M. Chambers, M. E. Stokes, P. H. Bos, A. Zask, Y. Zhang, M. Sanchez-Martin, M. A. Badgley, C. S. Huang, et al., Cell 2017, 168, 878–889.e29.

[15] M. P. Patricelli, M. R. Janes, L. S. Li, R. Hansen, U. Peters, L. V Kessler, Y. Chen, J. M. Kucharski, J. Feng, T. Ely, et al., Cancer Discov. 2016, 6, 316–329.

[16] M. R. Janes, J. Zhang, L. S. Li, R. Hansen, U. Peters, X. Guo, Y. Chen, A. Babbar, S. J. Firdaus, L. Darjania, et al., Cell 2018, 172, 578–589.

[17] E. J. Brown, M. W. Albers, T. Bum Shin, K. Ichikawa, C. T. Keith, W. S. Lane, S. L. Schreiber, Nature 1994, 369, 756–758.

[18] D. M. Sabatini, H. Erdjument-Bromage, M. Lui, P. Tempst, S. H. Snyder, Cell 1994, 78, 35–43.

[19] J. Krönke, N. D. Udeshi, A. Narla, P. Grauman, S. N. Hurst, M. Mcconkey, T. Svinkina, D. Heckl, E. Comer, X. Li, et al., Science 2014, 343, 301–305.

[20] Y. J. Li, H. X. Wang, H. F. Wu, Science 2014, 343, 305–309.

[21] I. M. Ahearn, F. D. Tsai, H. Court, M. Zhou, B. C. Jennings, M. Ahmed, N. Fehrenbacher, M. E. Linder, M. R. Philips, Mol. Cell 2011, 41, 173–185.

[22] J. Liu, J. D. Farmer, W. S. Lane, J. Friedman, I. Weissman, S. L. Schreiber, Cell 1991, 66, 807–815.

[23] K. H. Pua, D. T. Stiles, M. E. Sowa, G. L. Verdine, Cell Rep. 2017, 18, 432–442.

[24] A. C. Lai, M. Toure, D. Hellerschmied, J. Salami, S. Jaime-Figueroa, E. Ko, J. Hines, C. M. Crews, Angew. Chem. Int. Ed. 2016, 55, 807–810.

[25] T. Inoue, W. Do Heo, J. S. Grimley, T. J. Wandless, T. Meyer, Nat. Methods 2005, 2, 415–418.

[26] T. Bondeva, A. Balla, P. Várnai, T. Balla, Mol. Biol. Cell 2002, 13, 2323–2333.

[27] T. R. Brtva, J. K. Drugan, S. Ghosh, R. S. Terrell, S. Campbell-Burk, R. M. Bell, C. J. Der, J. Biol. Chem. 1995, 270, 9809–9812.

[28] T. W. Corson, N. Aberle, C. M. Crews, ACS Chem. Biol. 2010, 5, 63–77.

[29] S. L. Schreiber, Isr. J. Chem. 2018, 52–59.

