## Supplemental Information for "Bifunctional Small Molecule Ligands of K-Ras Induce Its Association with Immunophilin Proteins"

### Supporting Information

Supplementary Figures……………………………………………………………………………………….. 2
Methods……………………………………………………………………………………………………... 6
Materials
    List of Antibodies……………………………………………………………………………….. 11
    List of Buffer Composition………………………………………………………………………. 11
    List of Protein Sequences Used in This Study…………………………………………………. 12
Chemical Synthesis…………………………………………………………………………………….. 13

| Compound | Kd(FKBP12)/nM | Kd(CypA)/nM |
|---|---|---|
| **1** | 129 | – |
| **2** | 90 | – |
| **3** | 93 | – |
| **4** | 20.8 | – |
| **5** | 20 | – |
| **6** | 34.3 | – |
| **7** | – | 116 |
| **8** | – | 42.2 |
| **9** | – | 17.3 |
| **10** | – | 113 |
| **11** | 17.6 | – |
| **12** | – | 45.2 |

**Supplementary Figure 1.** Affinity of synthesized bifunctional ligands for FKBP12 or CypA determined by competition fluorescence anisotropy.

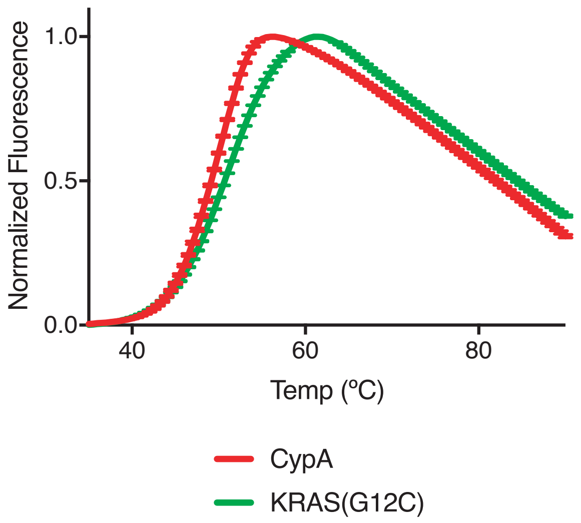


**Supplementary Figure 2.** Thermal denaturation curves of KRAS(G12C)•GDP and CypA.

**a**

| Drug (10 $\mu$M) | DMSO | **4** | **5** | **6** | ARS16 20 |
|---|---|---|---|---|---|
| pERK/ERK | 1.00 | 1.14 | 2.70 | 1.57 | 0.68 |
| pS6/S6 | 1.00 | 0.72 | 0.61 | 0.87 | 0.38 |

| Drug (10 $\mu$M) | DMSO | **7** | **8** | **9** | **10** | ARS16 20 |
|---|---|---|---|---|---|---|
| pERK/ERK | 1.00 | 1.01 | 1.06 | 1.74 | 0.84 | 0.40 |
| pS6/S6 | 1.00 | 0.08 | 0.59 | 0.32 | 0.86 | 0.36 |

**b**

| | **Empty Vector** | | **FKBP12** | | **Lyn-FKBP12** | |
|---|---|---|---|---|---|---|
| Drug (10 $\mu$M) | **–** | **5** | **–** | **5** | **–** | **5** |
| pERK/ERK | 1.00 | 0.93 | 1.29 | 1.06 | 0.94 | 0.74 |
| pS6/S6 | 1.00 | 0.92 | 1.07 | 0.85 | 0.89 | 0.70 |

| | **Empty Vector** | | **CypA** | | **Lyn-CypA** | |
|---|---|---|---|---|---|---|
| Drug (10 $\mu$M) | – | **9** | – | **9** | – | **9** |
| pERK/ERK | 1.00 | 0.99 | 1.24 | 1.02 | 1.04 | 0.48 |
| pS6/S6 | 1.00 | 0.66 | 1.03 | 0.78 | 0.85 | 0.52 |

**Supplementary Figure 3.** Quantification of phospho-ERK and phospho-S6 signals in Figure 4 by densitometry. Relative intensities were normalized to the respective DMSO-treated samples in **a** and empty vector-transfected, DMSO-treated samples in **b**.

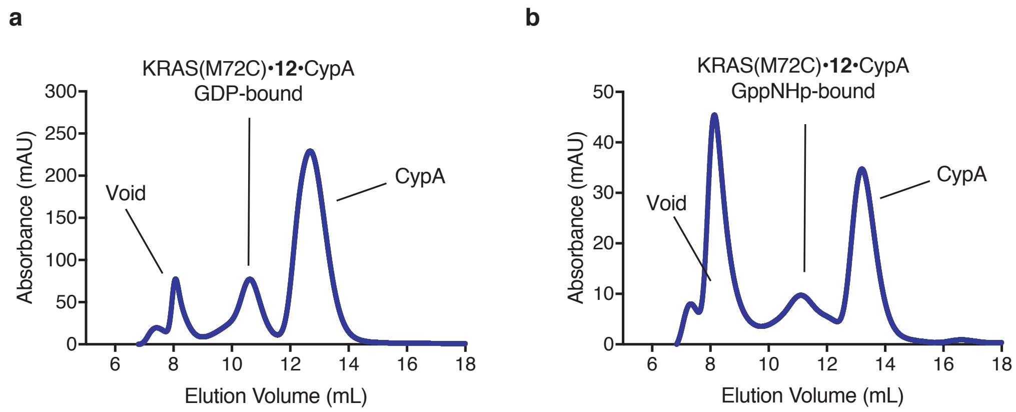


**Supplementary Figure 4.** Purification of K-Ras(M72C)•**12**•CypA (GDP-bound) and K-Ras(M72C)•**12**•CypA (GppNHp-bound).

**Expression and Purification of Recombinant FKBP12 and CypA**

DNA sequences encoding human FKBP12 and cyclophilin A were synthesized by Twist Biosciences and cloned into pET47b vector using standard molecular biology techniques (the sequences for these proteins are attached at the end of this document).
Protein expression was performed in BL21(DE3) *E. coli* strain. Briefly, chemically competent BL21(DE3) cells were transformed with pET47b-FKBP12 or pET47b-CypA and grown on LB agar plates containing 50 µg/mL kanamycin at 37 ºC. A single colony was used to inoculate a culture at 37 ºC, 220 rpm in terrific broth containing 50 µg/mL kanamycin. When the optical density reached 0.6, protein expression was induced by the addition of IPTG to 1 mM. After 2 h at 37 ºC, the cells were pelleted by centrifugation (6,500 x g, 10 min) and lysed in lysis buffer [20 mM Tris 8.0, 500 mM NaCl, 5 mM imidazole] with a high-pressure homogenizer (Microfluidics, Westwood, MA). The lysate was clarified by high-speed centrifugation (19,000 x g, 15 min) and the supernatant was used in subsequent purification by immobilized metal affinity chromatography (IMAC). His-tagged FKBP12 or CypA was captured by incubation with Co-TALON resin (Clonetech, Takara Bio USA, 4 mL slurry/liter culture) at 4 ºC for 1 h with constant end-to-end mixing. The loaded beads were then washed with lysis buffer (50 mL/liter culture) and the protein was eluted with elution buffer [20 mM Tris 8.0, 500 mM NaCl, 300 mM imidazole]. The His-tag was cleaved with His-tagged HRV 3C Protease (Clonetech, Takara Bio USA, 5 U/liter culture) at 4 ºC until LC-MS analysis of the reaction mixture indicated >95% cleavage. The reaction mixture was concentrated using an 10K MWCO centrifugal concentrator (Amicon-15, Millipore) to 20 mg/mL and purified by size exclusion chromatography on a Superdex 75 10/300 GL column (GE Healthcare Life Sciences) with SEC Buffer [20 mM HEPES 7.5, 150 mM NaCl, 1 mM $MgCl_2$]. Fractions containing pure FKBP12 or CypA protein were pooled and concentrated to 20 mg/mL and stored at –78 ºC. In our hands, this protocol gives a typical yield of 10-20 mg/liter culture for FKBP12, and 5-10 mg/liter culture for CypA.

**Expression and Purification of Recombinant KRAS CysLight G12C and M72C**

KRAS CysLight proteins were expressed and purified following previously reported protocols.[1,2]

**Expression and Purification of Biotinylated KRAS Proteins**

pET-Duet plasmids encoding BirA biotin-protein ligase and His-Avi-TEV-tagged KRAS proteins were constructed using standard molecular biology techniques. Biotinylated KRAS proteins were produced in BL21(DE3) *E. coli* strain. Briefly, chemically competent BL21(DE3) cells were transformed with the corresponding pET-Duet plasmid and grown on LB agar plates containing 100 µg/mL carbenicillin. A single colony was used to inoculate a culture at 37 ºC, 220 rpm in terrific broth containing 100 µg/mL carbenicillin. When the optical density reached 0.6, the culture temperature was reduced to 20 ºC, and protein expression was induced by the addition of IPTG to 1 mM and Biotin to 5 µM. After 16 h at 20 ºC, the cells were pelleted by centrifugation (6,500 x g, 10 min) and lysed in lysis buffer [20 mM Tris 8.0, 500 mM NaCl, 5 mM imidazole] with a high-pressure homogenizer (Microfluidics, Westwood, MA). The lysate was clarified by high-speed centrifugation (19,000 x g, 15 min) and the supernatant was used in subsequent purification by immobilized metal affinity chromatography (IMAC). His-Avi-TEV tagged K-Ras protein was captured with Co-TALON resin (Clonetech, Takara Bio USA, 2 mL slurry/liter culture) at 4 ºC for 1 h with constant end-to-end mixing. The loaded beads were then washed with lysis buffer (50 mL/liter culture) and the protein was eluted with elution buffer [20 mM Tris 8.0, 500 mM NaCl, 300 mM imidazole]. The protein was further purified with anion exchange chromatography (HiTrapQ column, GE Healthcare Life Sciences) using a salt gradient of 50 mM to 500 mM. Nucleotide loading was performed by mixing the ion exchange-purified protein with an excess of GDP (5 mg/liter culture) or GppNHp (5 mg/liter culture) and 5 mM EDTA at 23 ºC for 30 min. The reaction was stopped by the addition of $MgCl_2$ to 10 mM. For GppNHp, an additional calf intestine phosphatase treatment was performed as follows to ensure high homogeneity of the loaded nucleotide. The protein buffer was exchanged into Phosphatase Buffer [32 mM Tris 8.0, 200 mM ammonium sulfate, 0.1 mM $ZnCl_2$] with a HiTrap Desalting Column (GE Healthcare Life Sciences). To the buffer-exchanged protein solutions, GppNHp was added to 5 mg/mL, and Calf Intestine Phosphatase (NEB) was added to 10 U/mL. The reaction mixture was incubated on ice for 1 h, and $MgCl_2$ was added to a final concentration of 20 mM. After nucleotide loading, the protein was concentrated using an 10K MWCO centrifugal concentrator (Amicon-15, Millipore) to 20 mg/mL and purified by size exclusion chromatography on a Superdex 75 10/300 GL column (GE Healthcare Life Sciences). Fractions containing pure biotinylated Ras protein were pooled and concentrated to 20 mg/mL and stored at –78 ºC. In our hands, this protocol gives a typical yield of 5-15 mg/liter culture.

#### Expression and Purification of GST-Raf1-RBD and $Sos^{cat}$

The GST-tagged RBD domain of Raf1 (residues 1-149, GST-Raf1-RBD) was expressed and purified following published protocol. [3]
The catalytic domain of Sos (residues 466-1049, $Sos^{cat}$) was expressed and purified following published protocol.[4]

#### Other Proteins

GST-tagged full-length B-Raf protein (GST-BRAF) was purchased from MRC PPU (University of Dundee).

#### Determination of Compound Binding Affinity to FKBP12

Compound binding affinity was determined using a competition fluorescence polarization assay. A fluorescent tracer molecule based on SLF (FITC-SLF) was synthesized in house. The assay buffer was 20 mM HEPES 7.5, 0.01% Triton X-100. The Kd of the tracer molecule for FKBP12 was first determined by measuring fluorescence polarization (excitation 485 nm, emission 535 nm) at various protein concentrations and fitting the curve to a quadratic binding model. To measure compound binding affinity, mixtures with the following composition were prepared in duplicate in 96-well black opaque plates (Corning 3915): 5 nM fluorescent tracer, 10 nM FKBP12, 5% DMSO, 5 μM–0.08 nM of test compound, 200 μL total volume. Fluorescence polarization was measured on a TECAN Spark 20M plate reader (excitation 485 nm, emission 535 nm). Data were fitted to a three-parameter sigmoidal curve to derive $IC_{50}$ values. Kd of the compounds were calculated using a tool provided by Dr. Shaomeng Wang's lab (http://www.umich.edu/~shaomengwanglab/software/calc_ki/index.html).

#### Determination of Compound Binding Affinity to CypA

Compound binding affinity was determined using a competition fluorescence polarization assay. A fluorescent tracer molecule based on cyclosporin A (FITC-CsA) was synthesized in house. The assay buffer was 20 mM HEPES 7.5, 0.01% Triton-X-100. The Kd of the tracer molecule for CypA was first determined by measuring fluorescence polarization (excitation 485 nm, emission 535 nm) at various protein concentrations and fitting the curve to a quadratic binding model. To measure compound binding affinity, mixtures with the following composition were prepared in duplicate in 96-well black opaque plates (Corning 3915): 5 nM fluorescent tracer, 50 nM CypA, 5% DMSO, 5 μM–0.08 nM of test compound, 200 μL total volume. Fluorescence polarization was measured on a TECAN Spark 20M plate reader (excitation 485 nm, emission 535 nm). Data were fitted to a three-parameter sigmoidal curve to derive $IC_{50}$ values. Kd of the compounds were calculated using a tool provided by Dr. Shaomeng Wang's lab (http://www.umich.edu/~shaomengwanglab/software/calc_ki/index.html).

**Covalent Modification of K-Ras proteins by Small Molecules and Detection by Intact Protein Mass Spectrometry**

Test compounds were prepared as 100x stock solutions in DMSO. K-Ras proteins were diluted with Assay Buffer [20 mM HEPES 7.5, 150 mM NaCl, 1 mM $MgCl_2$] to 1 µM. In a typical reaction, 0.5 µL 100x compound stock was mixed with 50 µL diluted K-Ras protein, and the resulting mixture was incubated for the desired amount of time. Reactions were then stopped by addition of formic acid to a final concentration of 1% (v/v). The extent of modification was assessed by electrospray mass spectrometry using a Waters Acquity UPLC/ESI-TQD system equipped with a 2.1 x 50 mm Acquity UPLC BEH300 C4 column. The mobile phase was a linear gradient of 5-95% acetonitrile / water + 0.05% formic acid.

**Cell Culture**

NCI-H358 cells were obtained from ATCC and maintained in DMEM (Gibco) + 10% heat-inactivated FBS (Axenia Biologix) supplemented with penicillin and streptomycin (Gibco). When indicated, cells were treated with drugs at 60-80% confluency at a final DMSO concentration of 1%. At the end of treatment period, cells were placed on ice and washed once with PBS. The cells were scraped with a spatula, pelleted by centrifugation (500 x g, 5 min) and lysed in RIPA buffer supplemented with protease and phosphatase inhibitors (cOmplete and phosSTOP, Roche) on ice for 10 min. Lysates were clarified by high-speed centrifugation. Concentrations of lysates were determined with protein BCA assay (Thermo Fisher) and adjusted to 2 mg/mL with additional RIPA buffer. Samples were mixed with 5x SDS Loading Dye and heated at 95 ºC for 5 min.

**GST-FKBP12 and GST-CypA Pulldown**

Magnetic glutathione beads (Pierce, 25% slurry, 400 µL) were washed three times with 400 µL Co-IP Lysis Buffer. GST-FKBP12 (200 µg) or GST-CypA (200 µg) were then added to the washed beads as a solution in 400 µL Co-IP Lysis Buffer, and the mixture was incubated at 4 ºC for 1 h with constant end-to-end mixing. The supernatant was discarded, and the beads were washed three times with 400 µL Co-IP Lysis Buffer. In the last wash, beads were transferred into a new low-binding tube. Beads were resuspended with 400 µL Co-IP Lysis Buffer.
For each pulldown experiment, 50 µL GST-FKBP12 or GST-CypA beads slurry was captured on a magnetic stand. Supernatant was removed and 200 µL lysate (1 mg/mL) was added to the beads. The mixture was incubated at 23 ºC for 30 min with constant end-to-end mixing. Beads were washed twice with Co-IP Lysis Buffer (200 µL) and bound protein was eluted with 50 µL 1x LDS Loading Buffer at 95 ºC for 5 min.

**Gel Electrophoresis and Western Blot**

Unless otherwise noted, SDS-PAGE was run with Novex 4–12% Bis-Tris gel (Invitrogen) in MES running buffer (Invitrogen) at 200V for 40 min following the manufacturer's instructions. Protein bands were transferred onto 0.45-µm nitrocellulose membranes (Bio-Rad) using a wet-tank transfer apparatus (Bio-Rad Criterion Blotter) in 1x TOWBIN buffer with 10% methanol at 75V for 45 min. Membranes were blocked in 5% BSA–TBST for 1 h at 23 ºC. Primary antibody binding was performed with the indicated antibodies diluted in 5% BSA–TBST at 4 ºC for at least 16 h. After washing the membrane three times with TBST (5 min each wash), second antibodies (goat anti-rabbit IgG-IRDye 800 and goat anti-mouse IgG-IRDye 680, Li-COR) were added as solutions in 5% skim milk–TBST at the dilutions recommended by the manufacturer. Secondary antibody binding was allowed to proceed for 1 h at 23 ºC. The membrane was washed three times with TBST (5 min each wash) and imaged on a Li-COR Odyssey fluorescence imager.

**Differential Scanning Fluorimetry**

The protein of interest was diluted with SEC Buffer [20 mM HEPES 7.5, 150 mM NaCl, 1 mM $MgCl_2$] to 8 µM and mixed with a 100x DMSO solution of the ligand of interest. When no ligand was added, DMSO was used. SYPRO Orange Dye (Invitrogen) was added as a 500x solution in DMSO to a final nominal concentration of 5x. The resulting mixture was dispensed into wells of a white 96-well PCR plate in triplicate (25 µL/well). Fluorescence was measured at 0.5-ºC temperature intervals every 30 s from 25 ºC to 95 ºC on a Bio-Rad CFX96 qPCR system using the FRET setting. Each data set was normalized to the highest fluorescence and the normalized fluorescence reading was plotted against temperature in GraphPad Prism 6.0. Tm values were determined as the temperature(s) corresponding to the maximum(ma) of the first derivative of the curve.

**Thermal Denaturation of KRAS•CypA Complex**

40 µL KRAS(G12C)•**8**•CypA complex (1 mg/mL) was heated at the indicated temperature for 10 min, then was chilled on ice. The precipitated proteins were pelleted by centrifugation (21,000 x g, 10 min). The denaturation was performed in excess volume in order to prevent disturbing the pellet. Supernatants (8 µL) were mixed with 2 µL 5x SDS Loading Buffer and analyzed by SDS-PAGE.

**Sos- or EDTA-mediated Nucleotide Exchange Assay**

This assay was performed as previously reported[1,5–7] with slight modifications.
Mant-GDP (Invitrogen) was diluted to 1 µM with Nucleotide Exchange Assay Buffer. This is termed [Mant-GDP buffer] hereafter. Ras protein was diluted to 1.25 µM with Mant-GDP Buffer. 12 µL of this solution (triplicate for each condition) was added to wells of a black 384-well low-volume assay plate (Corning 4514). 3 µL of either Mant-GDP Buffer, 5 µM Sos in Mant-GDP Buffer, or 20 mM EDTA in Mant-GDP Buffer was added via a multichannel pipet rapidly to the wells. This should take less than15 s to finish. The plate was immediately placed in a TECAN Spark 20M plate reader, and fluorescence for Mant (excitation 360 nm, emission 440 nm) was read every 30 s over 1 h. Fluorescence intensity was normalized to values at time 0 and plotted again time.

**Time-Resolved Fluorescence Resonance Energy Transfer (TR-FRET) Assay**

TR-FRET assay was performed using recommended conditions from Cisbio with slight modifications. Biotinylated Ras proteins were diluted in TR-FRET buffer to 1.2 µM. GST-Raf1-RBD or GST-BRAF was in TR-FRET buffer to 400 nM. Anti-GST-Tb (Cisbo) was diluted to 500 ng/mL. Streptavidin-XL665 Cisbio was diluted to 4 µg/mL. For each replicate of each assay condition, 5 µL of diluted Ras protein and 5 µL diluted GST-Raf1-RBD or GST-BRAF was mixed in a well of a black low-volume 384-well plate (Corning 4514), and the mixture was incubated at 23 ºC for 1 h. 5 µL of Anti-GST-Tb and 5 µL of Streptavidin-XL665 were then added sequentially to each assay well, and the mixture was incubated at 23 ºC for an additional 1 h. Time-resolved fluorescence was read on a TECAN Spark 20M plate reader with the following parameters:
Lag time: 60 µs
Integration time: 500 µs
Read A: Excitation filter 320(25) nm, Emission filter 610(25) nm, Gain 130
Read B: Excitation filter 320(25) nm, Emission filter 665(8) nm, Gain 165
TR-FRET signal was calculated as the ratio fluorescence intensity [Read B]/[Read A]. The ratiometric signal was further normalized to a negative control containing no Ras protein.
Three replicates were performed for each assay condition.

**List of Antibodies**

| Target | Supplier | Identifier | Dilution |
|---|---|---|---|
| Pan-Ras | abcam | 108062 | 1:5000 |
| P-ERK [T202/Y204] | Cell Signaling Technology | 9101 | 1:1000 |
| Total ERK | Cell Signaling Technology | 4695 | 1:1000 |
| P-S6 [S240/S244] | Cell Signaling Technology | 5364 | 1:2000 |
| S6 | Cell Signaling Technology | 2217 | 1:1000 |
| FKBP12 | abcam | 58072 | 1:1000 |
| CypA | Cell Signaling Technology | 2175 | 1:1000 |

**List of Buffer Composition**

| | |
|---|---|
| Lysis Buffer | 20 mM Tris 8.0<br>500 mM NaCl<br>5 mM imidazole |
| Elution Buffer | 20 mM Tris 8.0<br>300 mM NaCl<br>300 mM imidazole |
| Phosphatase Buffer | 32 mM Tris 8.0<br>200 mM ammonium sulfate<br>0.1 mM $ZnCl_2$ |
| SEC Buffer | 20 mM HEPES 8.0<br>150 mM NaCl<br>1 mM $MgCl_2$ |
| Nucleotide Exchange Buffer | 20 mM HEPES 7.5<br>150 mM NaCl<br>1 mM $MgCl_2$<br>1 mM DTT |
| TR-FRET Buffer | 20 mM HEPES 7.5<br>150 mM NaCl<br>1 mM $MgCl_2$<br>0.05% Tween-20<br>0.1% BSA<br>0.5 mM DTT |
| Co-IP Lysis Buffer | 50 mM HEPES 7.5<br>120 mM NaCl<br>1 mM EDTA<br>1% IGEPAL CA 630 |

**List of Protein Sequences Used in This Study**

Texts in red indicate the affinity tags that were cleaved during purification.

**>FKBP12**
MAHHHHHHHHSAALEVLFQGPGYQGVQVETISPGDGRTFPKRGQTCVVHYTGMLEDGKKFDSSRDRNKPFKFMLGKQEVIR
GWEEGVAQMSVGQRAKLTISPDYAYGATGHPGIIPPHATLVFDVELLKLE

**>CypA**
MAHHHHHHHHSAALEVLFQGPDMVNPTVFFDIAVDGEPLGRVSFELFADKVPKTAENFRALSTGEKGFGYKGSCFHRIIPG
FMCQGGDFTRHNGTGGKSIYGEKFEDENFILKHTGPGILSMANAGPNTNGSQFFICTAKTEWLDGKHVVFGKVKEGMNIVE
AMERFGSRNGKTSKKITIADCGQLE

**>KRAS CysLight (G12C)**
MHHHHHHSSGRENLYFQGMTEYKLVVVGACGVGKSALTIQLIQNHFVDEYDPTIEDSYRKQVVIDGETSLLDILDTAGQEE
YSAMRDQYMRTGEGFLLVFAINNTKSFEDIHHYREQIKRVKDSEDVPMVLVGNKSDLPSRTVDTKQAQDLARSYGIPFIET
SAKTRQGVDDAFYTLVREIRKHKEK

**>KRAS CysLight (M72C)**
MHHHHHHSSGRENLYFQGMTEYKLVVVGAGGVGKSALTIQLIQNHFVDEYDPTIEDSYRKQVVIDGETSLLDILDTAGQEE
YSAMRDQYCRTGEGFLLVFAINNTKSFEDIHHYREQIKRVKDSEDVPMVLVGNKSDLPSRTVDTKQAQDLARSYGIPFIET
SAKTRQGVDDAFYTLVREIRKHKEK

**>His-Avi-TEV-KRAS (G12C)**
MGSSHHHHHHSGMSGLNDIFEAQKIEWHESSGENLYFQGMTEYKLVVVGACGVGKSALTIQLIQNHFVDEYDPTIEDSYRK
QVVIDGETSLLDILDTAGQEEYSAMRDQYMRTGEGFLLVFAINNTKSFEDIHHYREQIKRVKDSEDVPMVLVGNKSDLPSR
TVDTKQAQDLARSYGIPFIETSAKTRQGVDDAFYTLVREIRKHKEK

**>His-Avi-TEV-KRAS (M72C)**
MGSSHHHHHHSGMSGLNDIFEAQKIEWHESSGENLYFQGMTEYKLVVVGAGGVGKSALTIQLIQNHFVDEYDPTIEDSYRK
QVVIDGETSLLDILDTAGQEEYSAMRDQYCRTGEGFLLVFAINNTKSFEDIHHYREQIKRVKDSEDVPMVLVGNKSDLPSR
TVDTKQAQDLARSYGIPFIETSAKTRQGVDDAFYTLVREIRKHKEK

**>GST-Raf1-RBD**
MSPILGYWKIKGLVQPTRLLLEYLEEKYEEHLYERDEGDKWRNKKFELGLEFPNLPYYIDGDVKLTQSMAIIRYIADKHNM
LGGCPKERAEISMLEGAVLDIRYGVSRIAYSKDFETLKVDFLSKLPEMLKMFEDRLCHKTYLNGDHVTHPDFMLYDALDVV
LYMDPMCLDAFPKLVCFKKRIEAIPQIDKYLKSSKYIAWPLQGWQATFGGGDHPPKSDLVPRGSPIHIMEHIQGAWKTISN
GFGFKDAVFDGSSCISPTIVQQFGYQRRASDDGKLTDPSKTSNTIRVFLPNKQRTVVNVRNGMSLHDCLMKALKVRGLQPE
CCAVFRLLHEHKGKKARLDWNTDAASLIGEELQVDFLDHVPLTTHNFARKTFLKLGIHRD

**>GST-BRAF**
MSPILGYWKIKGLVQPTRLLLEYLEEKYEEHLYERDEGDKWRNKKFELGLEFPNLPYYIDGDVKLTQSMAIIRYIADKHNM
LGGCPKERAEISMLEGAVLDIRYGVSRIAYSKDFETLKVDFLSKLPEMLKMFEDRLCHKTYLNGDHVTHPDFMLYDALDVV
LYMDPMCLDAFPKLVCFKKRIEAIPQIDKYLKSSKYIAWPLQGWQATFGGGDHPPKSDLEVLFQGPLGSPNSRVDAALSGG
GGGGAEPGQALFNGDMEPEAGAGAGAAASSAADPAIPEEVWNIKQMIKLTQEHIEALLDKFGGEHNPPSIYLEAYEEYTSK
LDALQQREQQLLESLGNGTDFSVSSSASMDTVTSSSSSSLSVLPSSLSVFQNPTDVARSNPKSPQKPIVRVFLPNKQRTVV
PARCGVTVRDSLKKALMMRGLIPECCAVYRIQDGEKKPIGWDTDISWLTGEELHVEVLENVPLTTHNFVRKTFFTLAFCDF
CRKLLFQGFRCQTCGYKFHQRCSTEVPLMCVNYDQLDLLFVSKFFEHHPIPQEEASLAETALTSGSSPSAPASDSIGPQIL
TSPSPSKSIPIPQPFRPADEDHRNQFGQRDRSSSAPNVHINTIEPVNIDDLIRDQGFRGDGGSTTGLSATPPASLPGSLTN
VKALQKSPGPQRERKSSSSSEDRNRMKTLGRRDSSDDWEIPDGQITVGQRIGSGSFGTVYKGKWHGDVAVKMLNVTAPTPQ
QLQAFKNEVGVLRKTRHVNILLFMGYSTKPQLAIVTQWCEGSSLYHHLHIIETKFEMIKLIDIARQTAQGMDYLHAKSIIH
RDLKSNNIFLHEDLTVKIGDFGLATVKSRWSGSHQFEQLSGSILWMAPEVIRMQDKNPYSFQSDVYAFGIVLYELMTGQLP
YSNINNRDQIIFMVGRGYLSPDLSKVRSNCPKAMKRLMAECLKKKRDERPLFPQILASIELLARSLPKIHRSASEPSLNRA
GFQTEDFSLYACASPKTPIQAGGYGAFPVH

### Chemical Synthesis

### General Notes

Note on rotamers in $^1$H NMR data:

All of the SLF analogs and FK506 analogs synthesized here exist as a mixture of two amide rotamers in $CDCl_3$ or $CD_3OD$ (Mierke, D, F.; Schmieder, P.; Karuso, P.; Kessler, H. *Helv. Chim. Acta.* 1991, **74**, 1027–1047.). Due to extensive spectral overlap of the two, the coupling pattern of certain protons can be complicated even if they should display clear splitting patterns in theory. Sometimes, extensive spectral overlap prevents the identification of all peaks of the minor rotamer, and on occasion, of the major rotamer. In this document, only $^1$H NMR peaks of the major rotamer are reported in the best effort of resolving the peaks.

Cyclosporin analogs demonstrate more complicated conformational flexibility. In $CD_3OD$, most compounds exist as >6 conformational isomers (Ko, S. Y.; Dalvit, C. *Int. J. Pept. Protein Res. 1992*, **40,** 380–382.). In $CDCl_3$ the spectra are generally less complicated, and for certain compounds, only two conformational isomers are observed. For these compounds the $^1$H NMR spectra in $CDCl_3$ are resolvable, and peaks belonging to the major conformation are reported. For other compounds whose spectra are still too complicated to resolve, NMR data is not reported, and a note will be made in the experimental section.

Mini-workup

When a mini-workup (A/B) is indicated in the procedure, it was performed as follows: an aliquot (5 µL) of the reaction mixture was retrieved with a glass pipet and added to a plastic vial containing 0.2 mL organic solvent A and 0.2 mL aqueous solution B. The vial was shaken vigorously and allowed to stand until the two layers partitioned. The organic layer was then used for TLC or LC-MS analysis as specified in the procedure.

Monitoring Reaction Progress by LC-MS

When LC-MS analysis of the reaction mixture is indicated in the procedure, it was performed as follows. An aliquot (1 µL) of the reaction mixture (or the organic phase of a mini-workup mixture) was diluted with 100 µL 1:1 acetonitrile:water. 1 µL of the diluted solution was injected onto a Waters Acquity UPLC BEH C18 1.7 µm column and eluted with a linear gradient of 5–95% acetonitrile/water (+0.1% formic acid) over 3.0 min. Chromatograms were recorded with a UV detector set at 254 nm and a time-of-flight mass spectrometer (Waters Xevo G2-XS).

General Experiment Procedure

All reactions were performed in oven-dried glassware fitted with rubber septa under a positive pressure of argon, unless otherwise noted. Air- and moisture-sensitive liquids were transferred via syringe. Solutions were concentrated by rotary evaporation at or below 40 °C. Analytical thin-layer chromatography (TLC) was performed using glass plates pre-coated with silica gel (0.25-mm, 60-Å pore size, 230−400 mesh, Merck KGA) impregnated with a fluorescent indicator (254 nm). TLC plates were visualized by exposure to ultraviolet light (UV), then were stained by submersion in a 10% solution of phosphomolybdic acid (PMA) in ethanol or an acidic ethanolic solution of *p*-anisaldehyde,[1] followed by brief heating on a hot plate. Flash column chromatography was performed with Teledyne ISCO CombiFlash EZ Prep chromatography system, employing pre-packed silica gel

[1] This solution was prepared by sequential additions of concentrated sulfuric acid (5.0 mL), glacial acetic acid (1.5 mL) and *p*-anisaldehyde (3.7 mL) to absolute ethanol (135 mL) at 23 °C with efficient stirring.

cartridges (Teledyne ISCO RediSep).

Solvents and Reagents

Anhydrous solvents were purchased from Acros Organics. Except for those specified in the Starting Materials section, all chemical reagents were purchased from Sigma-Aldrich and AK Scientific. Commercial solvents and reagents were used as received.

Starting Materials

SLF was purchased from Cayman Chemical and/or synthesized following the synthetic route reported by Holt el al. Cyclosporin A and FK506 were purchased from LC Laboratories. tert-butyl 4-[7-bromo-6-chloro-2-[(3-ethoxy-3-oxo-propyl)amino]-8-fluoro-quinazolin-4-yl]piperazine-1-carboxylate (**S3**) was purchased in Pharmaron, Inc.

Instrumentation

Proton nuclear magnetic resonance ($^1$H NMR) spectra and carbon nuclear magnetic resonance ($^{13}$C NMR) spectra were recorded on Bruker AvanceIII HD 2-channel instrument (400 MHz/100 MHz) at 23 °C. Proton chemical shifts are expressed in parts per million (ppm, δ scale) and are referenced to residual protium in the NMR solvent ($CHCl_3$: δ 7.26, $D_2HCOD$: δ 3.31). Carbon chemical shifts are expressed in parts per million (ppm, δ scale) and are referenced to the carbon resonance of the NMR solvent ($CDCl_3$: δ 77.0, $CD_3OD$: δ 49.0). Data are represented as follows: chemical shift, multiplicity (s = singlet, d = doublet, t = triplet, q = quartet, dd = doublet of doublets, dt = doublet of triplets, m = multiplet, br = broad, app = apparent), integration, and coupling constant (*J*) in Hertz (Hz). High-resolution mass spectra were obtained using a Waters Xevo G2-XS time-of-flight mass spectrometer. Unless otherwise specified, diastereomeric ratios of products are reported as (major diastereomer) : (sum of minor diastereomers).

**Synthetic Procedure**

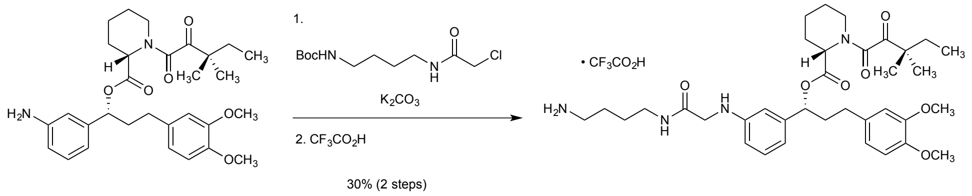


SLF S1

An oven-dried one-dram vial was charged with SLF (50 mg, 0.095 mmol), tert-butyl (4-(2-chloroacetamido)butyl)carbamate (37.8 mg, 0.143 mmol), potassium carbonate (19.7 mg, 0.143 mmol), DMF (0.19 mL) and a magnetic stir bar. The resulting mixture was warmed to 60 ºC and stirred for 24 h. The reaction mixture was partitioned between ethyl acetate (5 mL) and water (5 mL). The layers were separated, and the aqueous layer was extracted with ethyl acetate (2 x 5 mL). The combined organic layers were washed with saturated aqueous sodium chloride solution (15 mL). The washed solution was dried over sodium sulfate, and the dried solution was concentrated under reduced pressure. The residue was purified by column chromatography (20–50% ethyl acetate–hexanes) to afford the product as a colorless wax (21 mg).

Trifluoroacetic acid (0.5 mL) was added dropwise to a solution of the intermediate product (21 mg) in dichloromethane (0.5 mL) at 23 ºC. The resulting clear solution was allowed to stand at 23 ºC for 1 h. The solution was concentrated in vacuo to afford the product (**S1**) as a yellow solid (21 mg, 30% over 2 steps).

6:1 mixture of rotamers. Major rotamer is reported.

$^1$H NMR (400 MHz, Methanol-$d_4$) δ 7.20 (t, $J$ = 7.8 Hz, 1H), 6.88 (d, $J$ = 8.1 Hz, 1H), 6.85 – 6.78 (m, 1H), 6.78 – 6.70 (m, 2H), 6.66 – 6.59 (m, 2H), 5.68 (dd, $J$ = 8.7, 5.0 Hz, 1H), 5.22 (d, $J$ = 5.6 Hz, 1H), 3.83 (s, 3H), 3.82 (s, 3H), 3.79 (d, $J$ = 4.7 Hz, 2H), 3.42 (d, $J$ = 13.4 Hz, 1H), 3.31 – 3.14 (m, 3H), 2.91 (t, $J$ = 7.1 Hz, 2H), 2.72 – 2.50 (m, 3H), 2.37 (d, $J$ = 13.7 Hz, 1H), 2.32 – 2.18 (m, 1H), 2.12 – 2.00 (m, 1H), 1.85 – 1.43 (m, 8H), 1.43 – 1.29 (m, 2H), 1.25 (s, 3H), 1.24 (s, 3H), 0.91 (t, $J$ = 7.5 Hz, 3H).

HRMS (ESI): Calcd for $(C_{36}H_{52}N_4O_7 + H)^+$: 653.3916, Found: 653.3911.

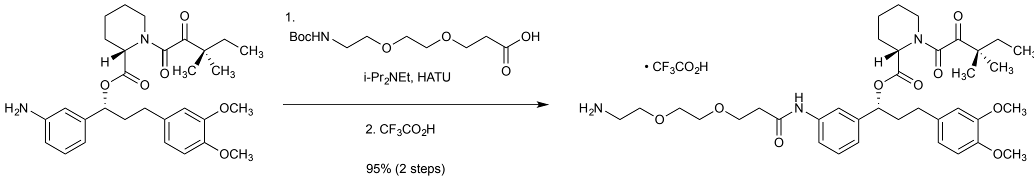


SLF S2

*N,N*-Diisopropylethylamine (16.6 µL, 0.10 mmol) and HATU (72 mg, 0.19 mmol) were added sequentially to a solution of SLF (50 mg, 0.10 mmol) and 3-[2-[2-(tert-butoxycarbonylamino)ethoxy]ethoxy]propanoic acid (40 mg, 0.14mmol) in 9:1 dichloromethane:DMF (0.2 mL). The resulting mixture was stirred at 23 ºC for 12 h, at which point LC-MS analysis showed full conversion to the desired product. The reaction mixture was partitioned between ethyl acetate (1 mL) and water (1 mL). The aqueous layer was extracted with ethyl acetate (3 x 1 mL). The combined organic layers were dried over sodium sulfate, then was concentrated under reduced pressure. The residue was purified by column chromatography (50–100% ethyl acetate–hexanes, 4-g CombiFlash column) to afford the product (**S2**) as a yellow oil.

The intermediate product (70 mg, 0.090 mmol) was dissolved in 1:1 dichloromethane:trifluoroacetic acid (1.0 mL). The resulting solution was allowed to stand at 23 ºC for 1 h, then was concentrated under reduced pressure to afford the product (**S2**) as a yellow foam (71 mg, 95%).

6:1 mixture of rotamers.

$^1$H NMR (400 MHz, Methanol-$d_4$) δ 7.70 (t, *J* = 1.9 Hz, 1H), 7.52 – 7.44 (m, 1H), 7.34 (t, *J* = 7.9 Hz, 1H), 7.14 (d, *J* = 8.0 Hz, 1H), 6.88 (d, *J* = 8.1 Hz, 1H), 6.81 (d, *J* = 2.1 Hz, 1H), 6.75 (dd, *J* = 8.2, 2.0 Hz, 1H), 5.73 (dd, *J* = 8.8, 5.0 Hz, 1H), 5.24 (d, *J* = 5.6 Hz, 1H), 3.86 (t, *J* = 6.5 Hz, 2H), 3.84 (s, 3H), 3.82 (s, 4H), 3.71 – 3.64 (m, 8H), 3.42 (d, *J* = 13.9 Hz, 1H), 3.26 (dd, *J* = 13.4, 3.3 Hz, 1H), 3.09 – 2.99 (m, 2H), 2.70 – 2.54 (m, 4H), 2.37 (d, *J* = 14.3 Hz, 1H), 2.34 – 2.22 (m, 1H), 2.15 – 2.03 (m, 1H), 1.84 – 1.59 (m, 5H), 1.59 – 1.29 (m, 2H), 1.25 (s, 3H), 1.24 (s, 3H), 0.90 (t, *J* = 7.5 Hz, 3H).

HRMS (ESI): Calcd for $(C_{37}H_{53}N_3O_9 + H)^+$: 684.3860, Found: 684.3851.

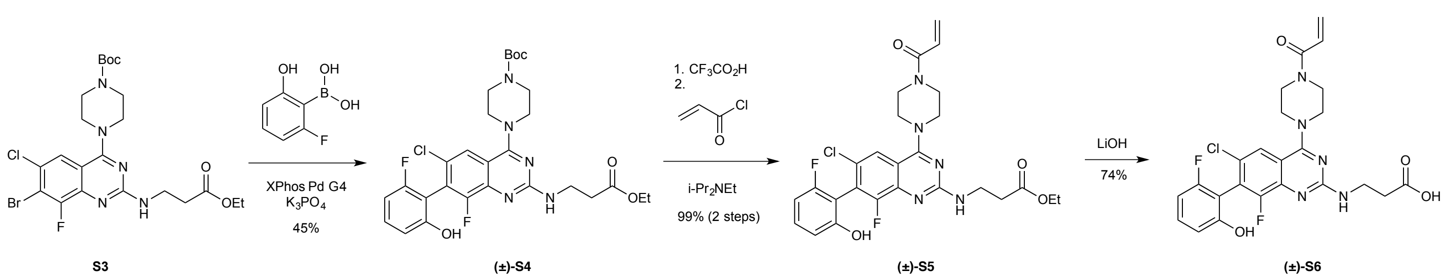


A 20-mL vial was charged with (2-fluoro-6-hydroxy-phenyl)boronic acid (278 mg, 1.78 mmol), tert-butyl 4-[7-bromo-6-chloro-2-[(3-ethoxy-3-oxo-propyl)amino]-8-fluoro-quinazolin-4-yl]piperazine-1-carboxylate (**S3**) (200 mg, 0.36 mmol), XPhos Pd G4 precatalyst (14.2 mg, 0.018 mmol), and 1:1 THF:water (4.0 mL). Argon was bubbled through the mixture for 5 min, then the vial was closed with a rubber septum fitted with a needle connected to an argon source. An 0.5 M aqueous solution of potassium phosphate (0.91 mL, 2.14 mmol) was added dropwise via syringe. After 16 h, both LC-MS and TLC analysis showed only ~50% conversion. Additional catalyst (14.2 mg) was added and the mixture was stirred at 50 ºC for another 2 h. However, no further progress was detected after this second portion of catalyst. The reaction mixture was partitioned between ethyl acetate (10 mL) and 10% citric acid (10mL). The layers were separated, and the aqueous layer was extracted with ethyl acetate (2 x 10 ml). The combined organic layers were dried over sodium sulfate, and the dried solution was concentrated. The residue was purified by column chromatography (20–50% ethyl acetate–hexanes, 4-g RediSep(R) Rf column, Teledyne ISCO, Lincoln, NE) to afford **(±)-S4** as a yellow powder (97 mg, 46%) and recovered bromide starting material (72 mg, 36%).

$^1$H NMR (400 MHz, Chloroform-*d*) δ 7.39 (s, 1H), 7.34 – 7.24 (m, 1H), 6.85 (d, *J* = 8.3 Hz, 1H), 6.70 (t, *J* = 8.5 Hz, 1H), 5.66 (s, 1H), 4.16 (q, *J* = 7.1 Hz, 2H), 3.74 (app q, *J* = 6.6 Hz, 2H), 3.71 – 3.51 (m, 8H), 2.65 (td, *J* = 6.5, 1.4 Hz, 2H), 1.50 (s, 9H), 1.26 (t, *J* = 7.1 Hz, 3H).

HRMS (ESI): Calcd for $(C_{28}H_{32}ClF_2N_5O_5 + H)^+$: 592.2138, Found: 592.2148.

**(±)-S4** (97 mg, 0.16 mmol) was dissolved in 1:1 trifluoroacetic acid (0.50 mL):dichloromethane (0.50 mL) and the resulting mixture was allowed to stand at 23 ºC for 1 h. The solution was then concentrated under reduced pressure to afford a yellow foam. Dichloromethane (1.45 mL) and *N,N*-diisopropylethylamine (43 µL, 0.25 mmol) were added sequentially to the foam. After stirring for 10 min, the material had fully dissolved. The solution was cooled to –78 ºC, and acryloyl chloride (14 µL, 0.170 mmol) was added dropwise via syringe. The resulting solution was warmed to 0 ºC, and the reaction progress was monitored by LC-MS. In 30 min, LC-MS analysis showed conversion to a single product with the desired m/z. The reaction mixture was partitioned between water (5 mL) and dichloromethane (5 mL). The layers were separated, and the aqueous layer was extracted with dichloromethane (2 x 5 mL). The combined organic layers were dried over sodium sulfate. The dried solution was filtered, and the filtrate was concentrated. The residue was purified by column chromatography (20–80% ethyl acetate–hexanes, 4-g RediSep(R) Rf column, Teledyne ISCO, Lincoln, NE) to afford **(±)-S5** as a yellow powder (100 mg, 99%).

$^1$H NMR (400 MHz, Methanol-$d_4$) δ 8.05 (s, 1H), 7.70 (dd, *J* = 13.0, 5.9 Hz, 1H), 7.45 – 7.29 (m, 3H), 6.82 (dt, *J* = 8.4, 0.8 Hz, 1H), 6.80 – 6.73 (m, 1H), 4.41 (s, 4H), 3.93 – 3.84 (m, 2H), 3.56 – 3.49 (m, 6H), 2.74 (t, *J* = 6.3 Hz, 2H), 1.20 (t, *J* = 7.0 Hz, 3H).

HRMS (ESI): Calcd for $(C_{26}H_{26}ClF_2N_5O_4 + H)^+$: 546.1719, Found: 546.1733.

Lithium hydroxide hydrate (1:1:1) (23 mg, 0.55 mmol) was added to a solution of **(±)-S5** (100 mg, 0.18 mmol) in 1:1 Water (1.10 mL):THF (1.10 mL). The resulting mixture was stirred at 23 °C, and the reaction progress was monitored by LC-MS. In 2 h, The ethyl ester had fully hydrolyzed. The volatile solvents were removed by rotary evaporation. The remaining aqueous suspension was acidified with 10% citric acid (2 mL), and then extracted with ethyl acetate (3 x 10 mL). The combined organic layers were dried over sodium sulfate, and the dried solution was concentrated to **(±)-S6** as a pale yellow powder (71 mg, 74%).

$^1$H NMR (400 MHz, Methanol-$d_4$) δ 7.83 (s, 1H), 7.32 (td, *J* = 8.3, 6.7 Hz, 1H), 6.91 – 6.77 (m, 2H), 6.73 (ddd, *J* = 9.2, 8.3, 1.0 Hz, 1H), 6.29 (dd, *J* = 16.8, 1.9 Hz, 1H), 5.82 (dd, *J* = 10.6, 1.9 Hz, 1H), 4.11 – 3.84 (m, 8H), 3.78 (t, *J* = 6.5 Hz, 2H), 2.68 (t, *J* = 6.5 Hz, 2H).

HRMS (ESI): Calcd for $(C_{24}H_{22}ClF_2N_5O_4 + H)^+$: 518.1396, Found: 518.1397.

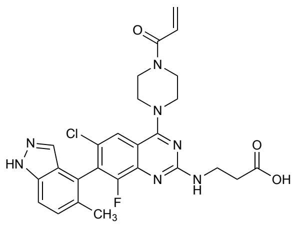


**(±)-S7**

**(±)-S7** was prepared using the same reaction sequence as for **(±)-S6**, except that 5-Methyl-1H-indazole-4-boronic acid was substituted for (2-fluoro-6-hydroxyphenyl)boronic acid.

**(±)-S7:** pale yellow powder.

$^{1}H$ NMR (400 MHz, DMSO-$d_6$) δ 13.13 (s, 1H), 7.81 (d, *J* = 1.4 Hz, 1H), 7.56 (d, *J* = 8.3 Hz, 2H), 7.37 (d, *J* = 8.6 Hz, 1H), 6.85 (dd, *J* = 16.7, 10.5 Hz, 1H), 6.17 (dd, *J* = 16.6, 2.4 Hz, 1H), 5.74 (dd, *J* = 10.4, 2.4 Hz, 1H), 3.88 – 3.61 (m, 8H), 3.61 – 3.48 (m, 4H), 2.16 (s, 3H).

HRMS (ESI): Calcd for $(C_{26}H_{25}ClFN_7O_3 + H)^+$: 538.1771, Found: 538.1766.

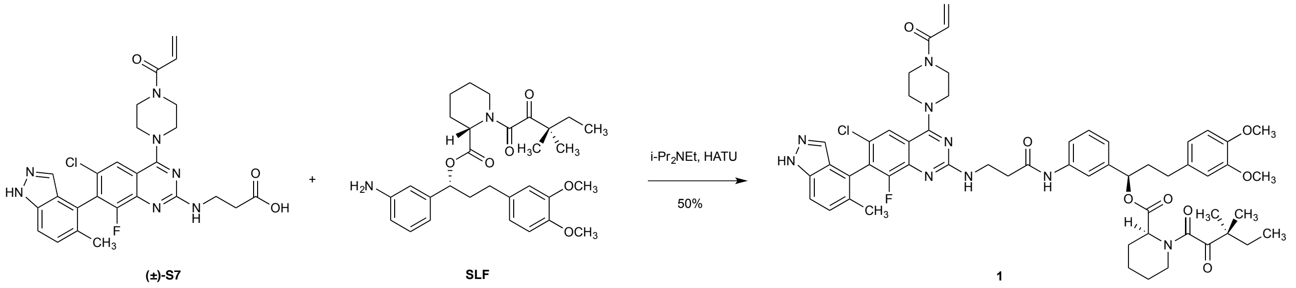


*N,N*-Diisopropylethylamine (4.2 µL, 0.024 mmol) and HATU (9.2 mg, 0.024 mmol) were added sequentially to a solution of **(±)-S7** (4.3 mg, 0.0080 mmol) and SLF (8.4 mg, 0.016 mmol) in DMF (0.20 mL) at 23 ºC. In 1 h, LC-MS analysis showed full consumption of the carboxylic acid component, with the formation of the desired mass. The reaction mixture was diluted with 50% acetonitrile–water to a volume of 4.7 mL, and the solution was purified by reverse-phase HPLC (Waters XBridge C18 column 5 µm particle size 30 x 250 mm, 5–95% acetonitrile–water + 0.1% formic acid, 40 min, 20 mL/min) to afford the product (**1**) as a white solid (4.2 mg, 50%).

**1** was a mixture of two diastereomers from the coupling of racemic **(±)-S7** and enantiomerically pure SLF. These two diastereomers were not separable with the HPLC condition described above, nor with other methods we have attempted.
$^{1}H$ NMR (400 MHz, Methanol-$d_4$) δ 8.14 (s, 1H), 7.90 (s, 1H), 7.73 – 7.52 (m, 2H), 7.53 – 7.40 (m, 1H), 7.39 – 7.23 (m, 2H), 7.13 – 7.01 (m, 2H), 6.93 – 6.78 (m, 3H), 6.78 – 6.65 (m, 1H), 6.29 (dd, *J* = 16.8, 2.0 Hz, 1H), 5.82 (dd, *J* = 10.6, 2.0 Hz, 1H), 5.75 (dd, *J* = 8.8, 4.8 Hz, 1H), 5.70 – 5.63 (m, 1H), 5.22 (s, 1H), 4.06 – 3.85 (m, 8H), 3.83 (s, 3H), 3.82 (s, 3H), 3.40 (d, *J* = 13.8 Hz, 1H), 3.29 – 3.13 (m, 1H), 2.78 – 2.53 (m, 4H), 2.42 – 2.29 (m, 3H), 2.24 (app d, *J* = 3.6 Hz, 3H), 2.22 – 1.98 (m, 1H), 1.87 – 1.61 (m, 6H), 1.40 – 1.29 (m, 2H), 1.23 (s, 6H), 0.89 (app q, *J* = 7.5 Hz, 3H).

HRMS (ESI): Calcd for $(C_{56}H_{63}ClFN_9O_8 + H)^+$: 1044.4552, Found: 1044.4580.

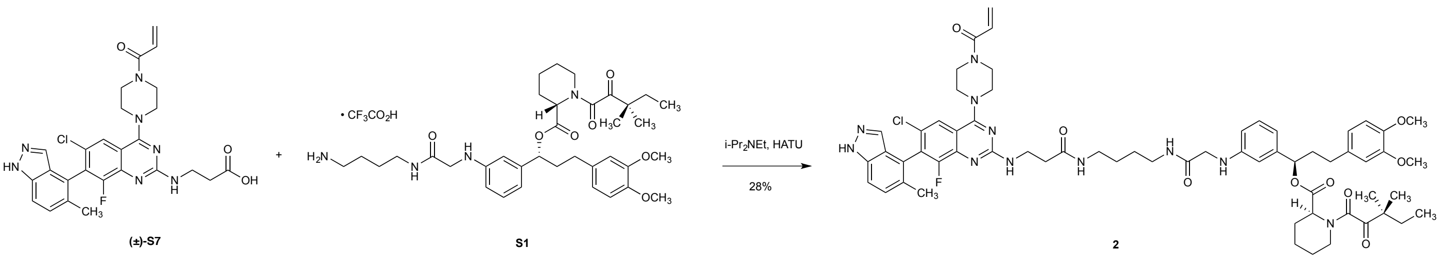


*N,N*-Diisopropylethylamine (4.9 µL, 0.028 mmol) and HATU (7.1 mg, 0.0186 mmol) were added sequentially to a mixture of **(±)-S7** (5.0 mg, 0.0093 mmol), **S1** (8.6 mg, 0.011 mmol), and DMF (0.2000 mL) at 23 ºC. In 1 h, LC-MS showed full consumption of the carboxylic acid starting material. The reaction mixture was diluted with

50% acetonitrile–water to a volume of 4.7 mL, and the solution was purified by reverse-phase HPLC (Waters XBridge C18 column 5 µm particle size 30 x 250 mm, 5–95% acetonitrile–water + 0.1% formic acid, 40 min, 20 mL/min) to afford the product (**2**) as a white solid (3.0 mg, 28%).

**2** was a mixture of two diastereomers from the coupling of racemic **(±)-S7** and enantiomerically pure SLF. These two diastereomers were not separable with the HPLC condition described above, nor with other methods we have attempted.

$^1$H NMR (400 MHz, Methanol-*d*$_4$) δ 7.88 (s, 1H), 7.77 (d, *J* = 9.3 Hz, 1H), 7.52 (d, *J* = 9.3 Hz, 1H), 7.17 – 7.09 (m, 1H), 6.90 – 6.81 (m, 2H), 6.81 – 6.74 (m, 2H), 6.74 – 6.64 (m, 2H), 6.61 – 6.49 (m, 2H), 6.29 (d, *J* = 16.2 Hz, 1H), 5.82 (dd, *J* = 10.6, 2.0 Hz, 1H), 5.73 – 5.58 (m, 1H), 5.25 – 5.17 (m, 1H), 4.00 – 3.83 (m, 8H), 3.81 (s, 3H), 3.80 (s, 3H), 3.74 – 3.68 (m, 2H), 3.47 – 3.41 (m, 1H), 3.23 – 3.06 (m, 2H), 3.06 – 2.90 (m, 4H), 2.68 – 2.48 (m, 4H), 2.39 – 2.29 (m, 2H), 2.23 (s, 3H), 2.10 – 1.97 (m, 1H), 1.83 – 1.59 (m, 6H), 1.50 – 1.35 (m, 6H), 1.23 (s, 3H), 1.22 (s, 3H), 0.89 (t, *J* = 7.1 Hz, 3H).

HRMS (ESI): Calcd for $(C_{62}H_{75}ClFN_{11}O_9 + H)^+$: 1172.5502, Found: 1172.5522.

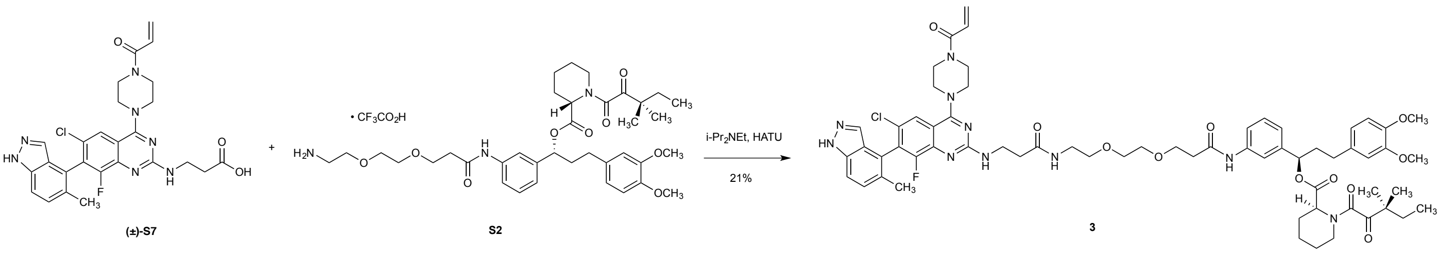


*N,N*-Diisopropylethylamine (4.9 µL, 0.028 mmol) and HATU (7.1 mg, 0.0186 mmol) were added sequentially to a mixture of **(±)-S7** (5.0 mg, 0.0093 mmol), **S2** (9.6 mg, 0.012 mmol), and DMF (0.2000 mL) at 23 ºC. In 1 h, LC-MS showed full consumption of the carboxylic acid starting material. The reaction mixture was diluted with 50% acetonitrile–water to a volume of 4.7 mL, and the solution was purified by reverse-phase HPLC (Waters XBridge C18 column 5 µm particle size 30 x 250 mm, 5–95% acetonitrile–water + 0.1% formic acid, 40 min, 20 mL/min) to afford the product (**3**) as a white solid (2.3 mg, 21%).

**3** was a mixture of two diastereomers from the coupling of racemic **(±)-S7** and enantiomerically pure SLF. These two diastereomers were not separable with the HPLC condition described above, nor with other methods we have attempted.

$^1$H NMR (400 MHz, Chloroform-*d*) δ 7.91 (s, 1H), 7.89 – 7.77 (m, 1H), 7.73 (d, *J* = 8.2 Hz, 1H), 7.69 – 7.41 (m, 2H), 7.07 – 6.97 (m, 1H), 6.84 – 6.75 (m, 2H), 6.75 – 6.63 (m, 3H), 6.64 – 6.56 (m, 1H), 6.44 (d, *J* = 16.8 Hz, 1H), 5.85 (d, *J* = 10.7 Hz, 1H), 5.83 – 5.70 (m, 1H), 5.36 – 5.26 (m, 1H), 4.31 – 4.16 (m, 4H), 4.07 – 3.92 (m, 4H), 3.87 (s, 3H), 3.86 (s, 3H), 3.81 (t, *J* = 5.1 Hz, 2H), 3.67 (s, 2H), 3.65 – 3.50 (m, 2H), 3.47 – 3.32 (m, 2H), 3.32 – 3.21 (m, 2H), 3.09 – 2.93 (m, 2H), 2.93 – 2.73 (m, 4H), 2.70 – 2.30 (m, 4H), 2.23 (s, 3H), 1.97 – 1.53 (m, 8H), 1.55 – 1.31 (m, 2H), 1.24 (s, 3H), 1.23 (s, 3H), 0.90 (t, *J* = 7.5 Hz, 3H).

HRMS (ESI): Calcd for $(C_{63}H_{76}ClFN_{10}O_{11} + H)^+$: 1203.5448, Found: 1203.5471.

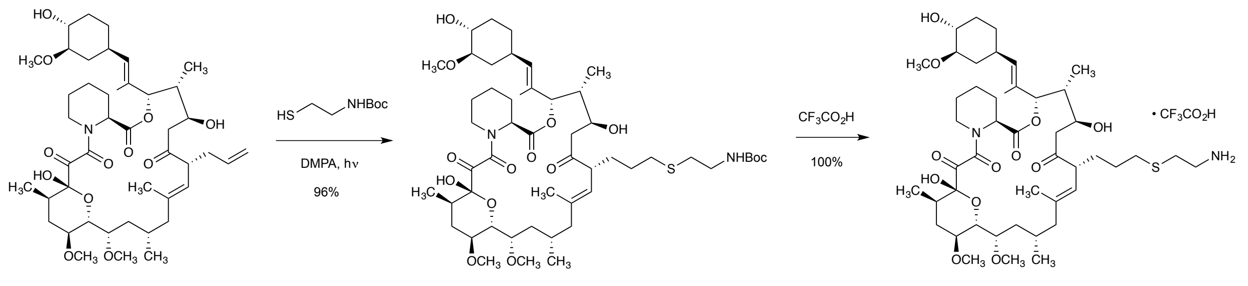


**S8** was prepared using a photo-catalyzed thiol-ene reaction reported by Guo et al. [8]

tert-Butyl N-(2-sulfanylethyl)carbamate (23 mg, 0.13 mmol) and dimethoxyphenylacetophenone (DMPA, 3.2 mg, 0.010 mmol) were added sequentially to a solution of FK506 (100 mg, 0.120 mmol) in dichloromethane (1.2 mL) at 23 ºC. After all reactants had dissolved, the vial was placed above a hand-held UV-light operating at 365 nm wavelength (the light was placed upside-down so that the contents of vial were directly irradiated). The irradiation was maintained for 15 min, at which point TLC analysis (100% ethyl acetate) showed full consumption of FK506 and formation of a slightly more polar product. The reaction solution was directly loaded onto a 12-g silica gel column. Purification by column chromatography (50–100% ethyl acetate–hexanes, 12-g RediSep(R) Rf column, Teledyne ISCO, Lincoln, NE) afforded the product (**S8**) as a white foam (117 mg, 96%).

Trifluoroacetic acid (0.5 mL) was added dropwise to a solution of **S8** (117 mg, 0.120 mmol) in dichloromethane (0.5 mL) at 0 ºC. The resulting mixture was stirred at 0 ºC for 1 h, at which point LC-MS analysis showed full deprotection of the Boc group. The reaction solution was concentrated under reduced pressure and the product was dried azeotropically by rotary evaporation of its suspension in toluene to afford the product (**S9**) as an off-white powder (119 mg, 100%).

3:2 mixture of rotamers.

$^{1}$H NMR (400 MHz, Methanol-$d_4$) δ 5.29 – 5.07 (m, 2H), 4.95 (d, $J$ = 11.7 Hz, 1H), 4.64 (s, 1H), 4.37 (d, $J$ = 13.4 Hz, 1H), 4.13 – 3.93 (m, 2H), 3.78 – 3.70 (m, 2H), 3.69 – 3.54 (m, 2H), 3.44 (s, 3H), 3.42 (s, 3H), 3.41 – 3.39 (m, 3H), 3.36 (s, 3H), 3.19 – 3.09 (m, 2H), 3.09 – 2.96 (m, 2H), 2.84 – 2.75 (m, 2H), 2.60 (t, $J$ = 6.9 Hz, 2H), 2.43 – 2.26 (m, 4H), 2.27 – 2.10 (m, 4H), 2.10 – 1.89 (m, 4H), 1.84 – 1.73 (m, 6H), 1.73 – 1.60 (m, 6H), 1.61 – 1.29 (m, 8H), 1.04 – 0.98 (m, 2H), 0.98 – 0.87 (m, 9H).

HRMS (ESI): Calcd for $(C_{46}H_{76}N_2O_{12}S + H)^+$: 881.5197, Found: 881.5207

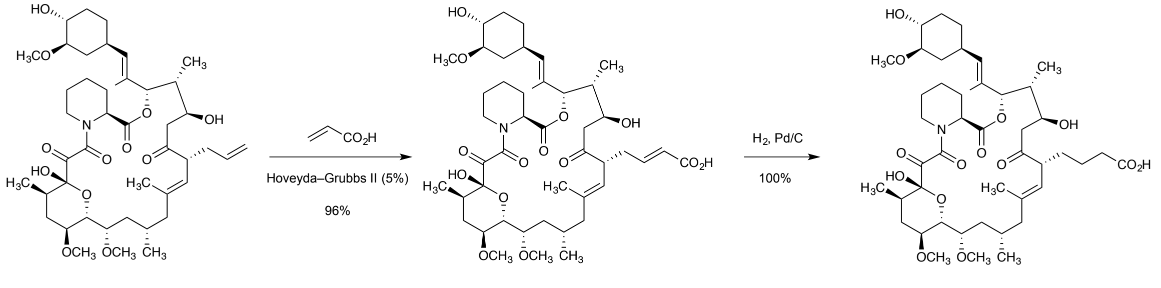


A flame-dried 10-mL microwave vial was flushed with dry argon, and then was charged with FK506 (100 mg, 0.120 mmol), DCE (1.20 mL), and a magnetic stir bar. Acrylic acid (170 mg, 2.49 mmol) and Grubbs Catalyst

2nd Gen (5.3 mg, 0.010 mmol) were added sequentially. The vial was flushed with argon again and sealed with a rubber cap. The reaction mixture was heated at 85 ºC for 1 h in a CEM Discover SP microwave reactor. After cooling to 23 ºC, TLC analysis (100% ethyl acetate) of the reaction mixture showed full disappearance of the starting material and formation of a highly polar new spot. The reaction solution was concentrated in vacuo. The residue was purified by column chromatography (20–100% ethyl acetate–hexanes, 12-g RediSep(R) Rf column, Teledyne ISCO, Lincoln, NE) to afford the product (**S10**) as a yellow powder (101 mg, 96%).

10 wt% Palladium on carbon (13 mg, 0.010 mmol) was added to a solution of **S10** (50 mg, 0.060 mmol) in methanol (5 mL) at 23 ºC under an atmosphere of argon. The reaction flask was evacuated until effervescence occurred, then flushed with hydrogen gas. The process was repeated three times. The resulting suspension was stirred at 23 ºC for 16 h under an atmosphere of hydrogen. The reaction flask was purged with argon, and the reaction suspension was filtered through a pad of Celite. The filter cake was rinsed with ethyl acetate (20 mL). The combined filtrate was concentrated in vacuo, and the residue was purified by column chromatography (0–10% methanol–dichloromethane + 0.1% acetic acid) to afford the product (**S11**) as a white solid (50 mg, 100%).

3:2 mixture of rotamers.

$^1$H NMR (400 MHz, Chloroform-*d*) δ 5.35 – 5.31 (m, 1H), 5.13 – 5.07 (m, 1H), 5.03 (d, *J* = 10.3 Hz, 1H), 4.61 (d, *J* = 5.4 Hz, 1H), 4.43 (d, *J* = 13.8 Hz, 1H), 4.01 – 3.89 (m, 1H), 3.70 (d, *J* = 9.6 Hz, 1H), 3.64 – 3.52 (m, 1H), 3.42 (s, 3H), 3.40 (s, 3H), 3.43 – 3.36 (m, 3H), 3.31 (s, 3H), 3.10 – 2.93 (m, 3H), 2.79 (dd, *J* = 15.9, 3.0 Hz, 1H), 2.45 – 2.25 (m, 5H), 2.26 – 2.10 (m, 3H), 2.09 – 1.88 (m, 6H), 1.89 – 1.71 (m, 6H), 1.60 (s, 6H), 1.60 – 1.35 (m, 8H), 1.13 – 1.03 (m, 2H), 1.00 (d, *J* = 6.3 Hz, 3H), 0.94 (d, *J* = 6.4 Hz, 3H), 0.88 (d, *J* = 7.1 Hz, 3H).

HRMS (ESI): Calcd for $(C_{45}H_{71}NO_{14} - H)^-$: 848.4796, Found: 848.4809

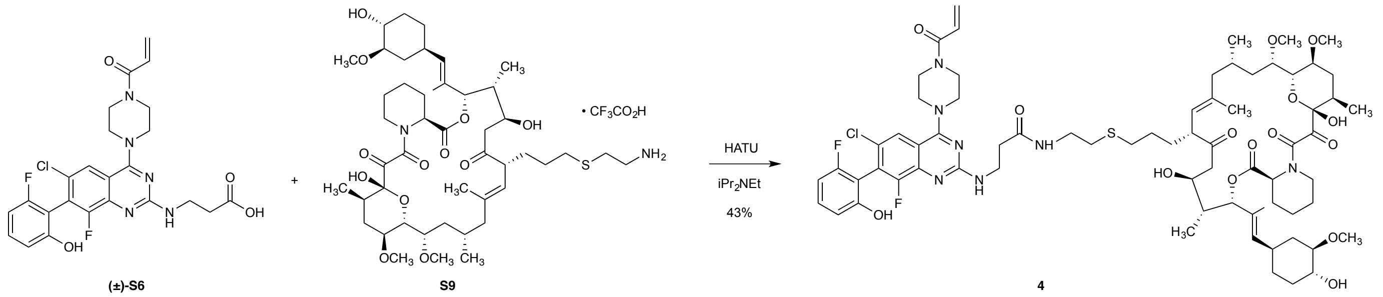


A 1-dram vial was charged with (±)-**S6** (5.0 mg, 0.010 mmol), **S9** (9.6 mg, 0.010 mmol), and DMF (0.30 mL). *N,N*-Diisopropylethylamine (5.0 µL, 0.030 mmol) and HATU (4.4 mg, 0.012 mmol) were added sequentially to the solution at 23 °C, and the resulting mixture was stirred at 23 °C while the reaction progress was monitored by LC-MS. In 15 min, LC-MS showed full consumption of the amine starting material and formation of a new product with the expected m/z. The reaction mixture was diluted with 50% acetonitrile–water to a volume of 4.7 mL, and the solution was filtered through a 0.45 µM PTFE syringe filter. The filtrate was purified by reverse-phase HPLC (Waters XBridge C18 column 5 µm particle size 30 x 250 mm, 5–95% acetonitrile–water + 0.1% formic acid, 40 min, 20 mL/min) to afford the product (**4**) as a white solid (5.7 mg, 43%).

3:2 mixture of rotamers.

$^1$H NMR (400 MHz, Chloroform-*d*) δ 8.28 (s, 1H), 7.77 – 7.62 (m, 1H), 6.90 – 6.72 (m, 2H), 6.64 (dd, *J* = 16.9, 10.7 Hz, 1H), 6.39 (d, *J* = 16.8 Hz, 1H), 5.80 (d, *J* = 10.3 Hz, 1H), 5.35 (d, *J* = 3.5 Hz, 1H), 5.11 – 4.96 (m, 2H), 4.61 – 4.32 (m, 2H), 4.07 – 3.73 (m, 6H), 3.72 – 3.53 (m, 2H), 3.42 (s, 3H), 3.42 (s, 3H), 3.41 (s, 3H), 3.40 – 3.37 (m, 3H), 3.17 – 2.86 (m, 6H), 2.68 – 2.52 (m, 4H), 2.49 – 2.24 (m, 6H), 2.24 – 1.86 (m, 8H), 1.86 – 1.70 (m, 6H), 1.70 – 1.61 (m, 6H), 1.61 – 1.24 (m, 8H), 1.14 – 1.05 (m, 2H), 1.05 – 0.81 (m, 9H).

HRMS (ESI): Calcd for $(C_{70}H_{96}ClF_2N_7O_{15}S + H)^+$: 1380.6420, Found: 1380.6404

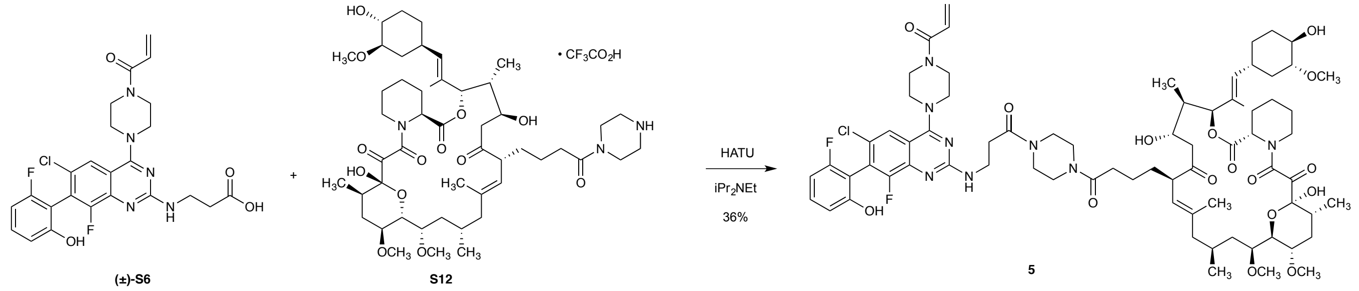


A 1-dram vial was charged with **(±)-S6** (5.5 mg, 0.011 mmol), **S12** (10.0 mg, 0.010 mmol), and DMF (0.30 mL). *N,N*-Diisopropylethylamine (5.0 µL, 0.030 mmol) and HATU (7.6 mg, 0.019 mmol) were added sequentially to the solution at 23 °C, and the resulting mixture was stirred at 23 °C while the reaction progress was monitored by LC-MS. In 15 min, LC-MS showed full consumption of the amine starting material and formation of a new product with the expected m/z. The reaction mixture was diluted with 50% acetonitrile–water to a volume of 4.2 mL, and the solution was filtered through a 0.45 µM PTFE syringe filter. The filtrate was purified by reverse-phase HPLC (Waters XBridge C18 column 5 µm particle size 30 x 250 mm, 5–95% acetonitrile–water + 0.1% formic acid, 40 min, 20 mL/min) to afford the product (**5**) as a white solid (4.9 mg, 36%).

3:2 mixture of rotamers.

$^1$H NMR (400 MHz, Methanol-*d*$_4$) δ 8.09 (s, 1H), 7.46 – 7.29 (m, 1H), 6.89 – 6.71 (m, 3H), 6.33 (d, *J* = 16.6 Hz, 1H), 5.85 (d, *J* = 10.6 Hz, 1H), 5.33 – 5.09 (m, 3H), 4.34 (br s, 4H), 4.06 – 3.83 (m, 4H), 3.77 – 3.52 (m, 10H), 3.43 (s, 3H), 3.42 (s, 3H), 3.41 – 3.38 (m, 2H), 3.35 (s, 3H), 3.09 – 2.94 (m, 2H), 2.94 – 2.75 (m, 4H), 2.50 – 2.27 (m, 6H), 2.27 – 1.87 (m, 6H), 1.88 – 1.72 (m, 4H), 1.72 – 1.52 (m, 6H), 1.52 – 1.28 (m, 8H), 1.15 – 1.02 (m, 2H), 1.02 – 0.86 (m, 9H).

HRMS (ESI): Calcd for $(C_{73}H_{99}ClF_2N_8O_{16} + H)^+$: 1417.6914, Found: 1417.6904

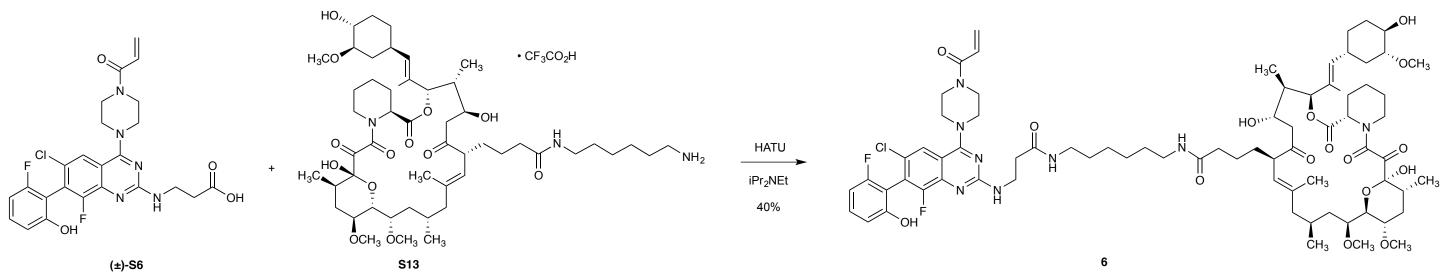

A 1-dram vial was charged with **(±)-S6** (5.5 mg, 0.011 mmol), **S12** (10.0 mg, 0.010 mmol), and DMF (0.30 mL). *N,N*-Diisopropylethylamine (5.0 μL, 0.030 mmol) and HATU (7.6 mg, 0.019 mmol) were added sequentially to the solution at 23 °C, and the resulting mixture was stirred at 23 °C while the reaction progress was monitored by LC-MS. In 15 min, LC-MS showed full consumption of the amine starting material and formation of a new product with the expected m/z. The reaction mixture was diluted with 50% acetonitrile–water to a volume of 4.2 mL, and the solution was filtered through a 0.45 μM PTFE syringe filter. The filtrate was purified by reverse-phase HPLC (Waters XBridge C18 column 5 μm particle size 30 x 250 mm, 5–95% acetonitrile–water + 0.1% formic acid, 40 min, 20 mL/min) to afford the product (**6**) as a white solid (5.5 mg, 40%).

3:2 mixture of rotamers.

$^1$H NMR (400 MHz, Methanol-*d*$_4$) δ 7.77 (s, 1H), 7.35 – 7.26 (m, 1H), 6.89 – 6.68 (m, 3H), 6.28 (d, *J* = 16.1 Hz, 1H), 5.82 (d, *J* = 12.3 Hz, 1H), 5.28 – 5.19 (m, 1H), 5.18 – 4.99 (m, 2H), 4.60 (s, 2H), 4.40 – 4.26 (m, 1H), 4.17 – 4.06 (m, 1H), 3.95 – 3.83 (m, 8H), 3.84 – 3.74 (m, 2H), 3.56 – 3.47 (m, 2H), 3.44 (s, 3H), 3.43 (s, 3H), 3.40 (s, 3H), 3.39 – 3.36 (m, 2H), 3.20 – 3.02 (m, 4H), 2.61 – 2.50 (m, 2H), 2.44 – 2.29 (m, 4H), 2.25 – 2.09 (m, 4H), 2.09 – 1.89 (m, 4H), 1.89 – 1.65 (m, 6H), 1.65 – 1.33 (m, 10H), 1.31 – 1.23 (m, 8H), 1.12 – 1.02 (m, 2H), 1.03 – 0.78 (m, 9H).

HRMS (ESI): Calcd for $(C_{75}H_{105}ClF_2N_8O_{16} + H)^+$: 1447.7383, Found: 1447.7435

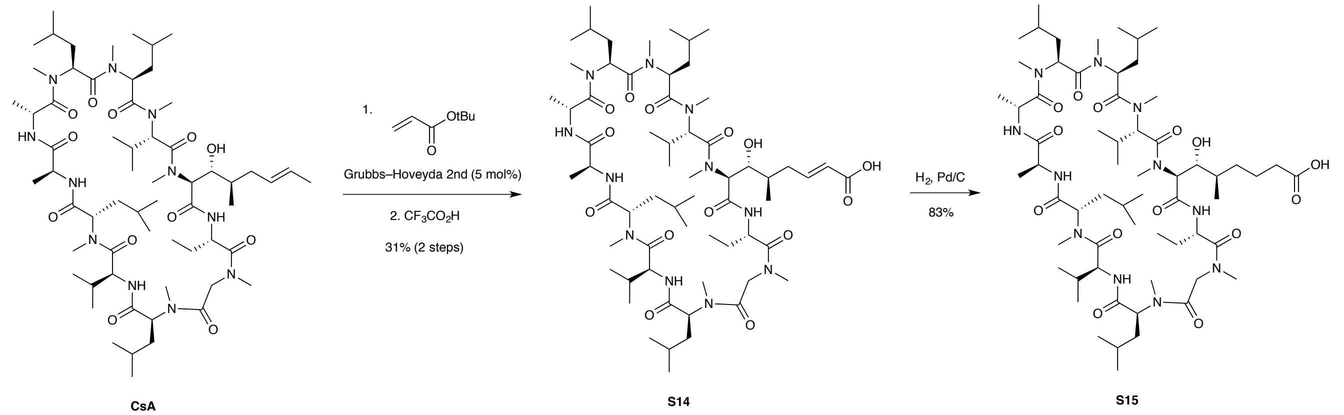


A flame-dried 10-mL microwave vial was flushed with dry argon, and then was charged with cyclosporin A (100 mg, 0.083 mmol), 1,2-dichloroethane (1.24 mL), and a magnetic stir bar. tert-Butyl acrylate (0.24 mL, 1.66 mmol) and Grubbs–Hoveyda Catalyst 2nd Gen (3.5 mg, 0.042 mmol) were added sequentially. The vial was flushed with argon again and sealed with a rubber cap. The reaction mixture was heated at 70 ºC for 1 h in a CEM Discover SP microwave reactor with constant stirring. After cooling to 23 ºC, LC-MS analysis showed ~50% conversion to the desired product mass. The vial was returned to the microwave reactor and heated for an additional 3 h at 70 °C. The reaction mixture was cooled to 23 ºC and directly loaded onto a silica gel cartridge (~4 g). Purification by column chromatography (0–20% methanol–dichloromethane, 4-g RediSep Rf Column, Teledyne ISCO, Lincoln, NE) afforded a brown solid. The purity of this material was ~80% by $^1$H NMR analysis.

Trifluoroacetic acid (0.5 mL) was added to a solution of the product from the cross-metathesis reaction in dichloromethane (0.5 mL). In 1 h, LC-MS analysis showed full consumption of the tert-butyl ester starting material (m/z = 1288). The reaction mixture was concentrated under vacuum. The residue was diluted with 50% acetonitrile–water to a volume of 4.0 mL, and the solution was filtered through a 0.45 μM PTFE syringe filter. The filtrate was purified by reverse-phase HPLC (Waters XBridge C18 column 5 μm particle size 30 x 250 mm,

5–95% acetonitrile–water + 0.1% formic acid, 40 min, 20 mL/min) to afford the product (**S14**) as a white solid (31 mg, 31% over 2 steps).

A 100-mL round bottom flask was charged with **S14** (412 mg, 0.33 mmol), 1:1 Ethyl acetate (6.6 mL):methanol (6.6 mL) and a magnetic stir bar. Argon was bubbled through the solution for 5 min, then Palladium on carbon (10 wt%, 71 mg, 0.033 mmol) was added. The vessel was fitted with a rubber septum and a hydrogen balloon was attached via a 19-gauge needle. An additional needle was attached to allow a gentle flow of hydrogen to bubble through the solution at a continuous rate. At 3 h, LC-MS could no longer detect any starting material. The hydrogen balloon was switched to one filled with argon, and bubbling was continued for 5 min. The reaction mixture was then filtered through a tightly packed plug of Celite (~2g). Concentration of the filtrate afforded a colorless glass. The material was purified by reverse-phase HPLC in multiple batches with the following procedure. The residue was divided into batches and dissolved in 50% methanol–water (100 mg in 5 mL, 150 mg in 8 mL, then 150 mg in 8 mL), and the solutions were filtered through a 0.45 μM PTFE syringe filter. The filtrate was purified by reverse-phase HPLC (Waters XBridge C18 column 5 μm particle size 30 x 250 mm, 50–95% acetonitrile–water + 0.1% formic acid, 40 min, 20 mL/min) in batches, and the product-containing fractions were pooled to afford the product (**S15**) as a white solid (343 mg, 83%).

$^1$H NMR (400 MHz, Chloroform-*d*) δ 7.99 (d, *J* = 9.2 Hz, 1H), 7.63 (d, *J* = 7.5 Hz, 1H), 5.68 (dd, *J* = 11.0, 4.1 Hz, 1H), 5.39 (d, *J* = 7.3 Hz, 1H), 5.31 – 5.23 (m, 1H), 5.17 – 5.05 (m, 3H), 5.00 (q, *J* = 7.5 Hz, 1H), 4.88 – 4.81 (m, 1H), 4.72 (d, *J* = 14.3 Hz, 1H), 4.63 (t, *J* = 8.8 Hz, 1H), 4.51 (dt, *J* = 14.6, 7.5 Hz, 1H), 3.87 (t, *J* = 6.3 Hz, 1H), 3.42 (s, 3H), 3.40 – 3.35 (m, 4H), 3.30 (d, *J* = 11.8 Hz, 1H), 3.23 (s, 3H), 3.19 (s, 3H), 3.18 (d, *J* = 17.2 Hz, 1H), 3.09 (s, 3H), 2.87 (d, *J* = 17.6 Hz, 1H), 2.78 – 2.68 (m, 1H), 2.69 (s, 3H), 2.67 (s, 3H), 2.48 – 2.30 (m, 2H), 2.23 (t, *J* = 6.9 Hz, 2H), 2.22 – 1.87 (m, 5H), 1.80 – 1.37 (m, 13H), 1.34 (d, *J* = 7.3 Hz, 3H), 1.26 (d, *J* = 6.9 Hz, 3H), 1.24 – 1.12 (m, 3H), 1.07 – 0.79 (m, 39H, 13 methyl doublets).

HRMS (ESI): Calcd for $(C_{62}H_{111}N_{11}O_{14} – H)^–$: 1232.8234, Found: 1232.8215

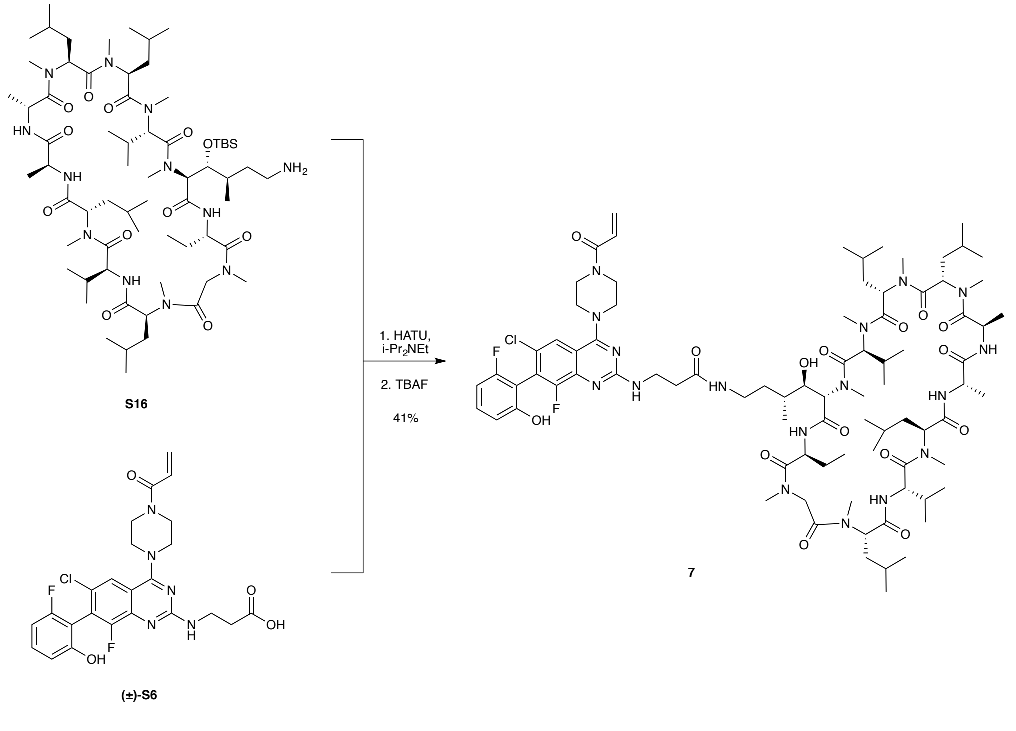

A 1-dram vial was charged with **S16** (7.0 mg, 0.0054 mmol), **(±)-S6** (3.3 mg, 0.0064 mmol) and a magnetic stir bar. DMF (0.10 mL) and *N,N*-Diisopropylethylamine (4.7 µL, 0.027 mmol) were added sequentially via pipette. HATU (2.5 mg, 0.0064 mmol) was added as a freshly prepared 10% w/v DMF solution via pipette. The resulting solution was stirred at 23 ºC and the reaction progress was monitored by LC-MS. In 12 h, LC-MS analysis showed full consumption of the amine starting material and formation of a new species. The reaction mixture was diluted with 0.1 mL THF, and a 1.0 M solution of tetra-n-butylammonium fluoride in THF (27 µL, 0.027 mmol) was added dropwise via syringe. In 2 h, LC-MS showed full deprotection of the TBS group. The residue was diluted with 50% acetonitrile–water to a volume of 3.9 mL, and the solution was filtered through a 0.45 µM PTFE syringe filter. The filtrate was purified by reverse-phase HPLC (Waters XBridge C18 column 5 µm particle size 30 x 250 mm, 50–95% acetonitrile–water + 0.1% formic acid, 40 min, 20 mL/min) to afford the product (**7**) as a white solid (3.7 mg, 41%).

HRMS (ESI): Calcd for $(C_{84}H_{130}ClF_2N_{17}O_{15} - H)^-$: 1688.9511, Found: 1688.9529.

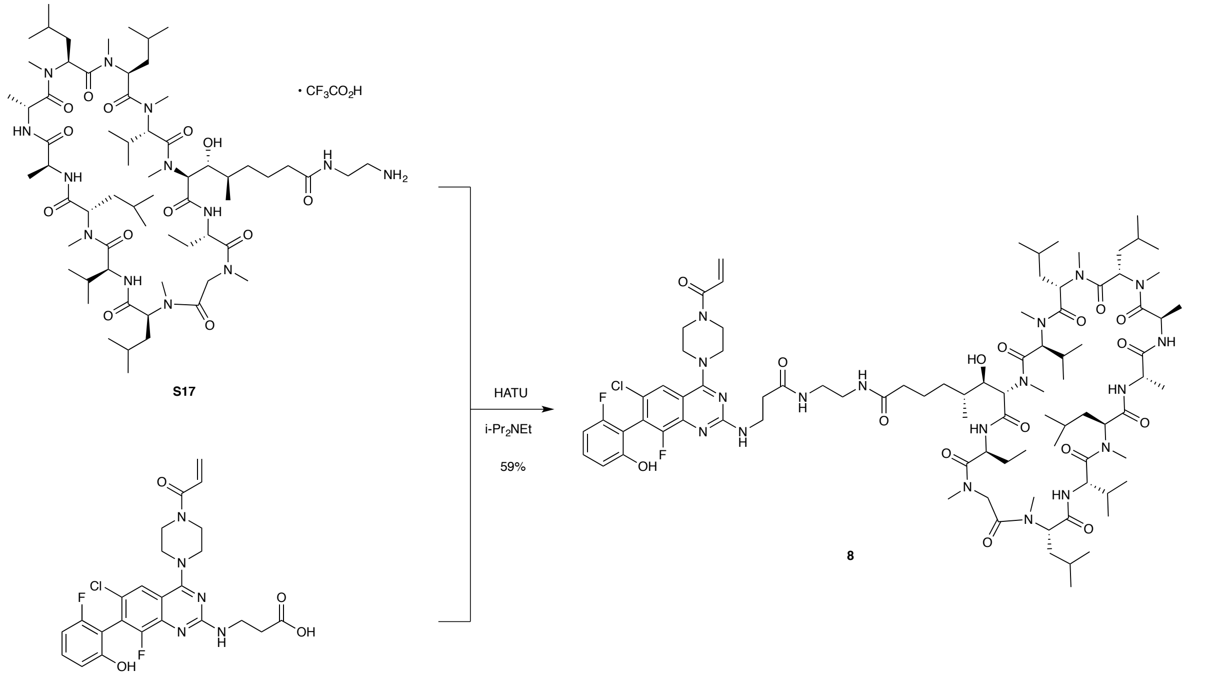


An oven dried one-dram vial was charged with **S17** (10 mg, 0.076 mmol), **(±)-S6** (4.3 mg, 0.083 mmol), DMF (0.10 mL), and a magnetic stir bar. *N,N*-Diisopropylethylamine (4.0 µL, 0.023 mmol) and HATU (4.3 mg, 0.011 mmol) were added sequentially to the reaction mixture at 23 ºC. In 40 min, LC-MS analysis showed 100% conversion. The reaction mixture was diluted with 50% acetonitrile–water to a volume of 5.0 mL, and the solution was filtered through a 0.45 µM PTFE syringe filter. The filtrate was purified by reverse-phase HPLC (Waters XBridge C18 column 5 µm particle size 30 x 250 mm, 5–95% acetonitrile–water + 0.1% formic acid, 40 min, 20 mL/min) to afford the product (**8**) as a white powder (7.9 mg, 58%).

HRMS (ESI): Calcd for $(C_{88}H_{137}ClF_2N_{18}O_{16} + 2H)^{2+}$: 888.5136, Found: 888.5137.

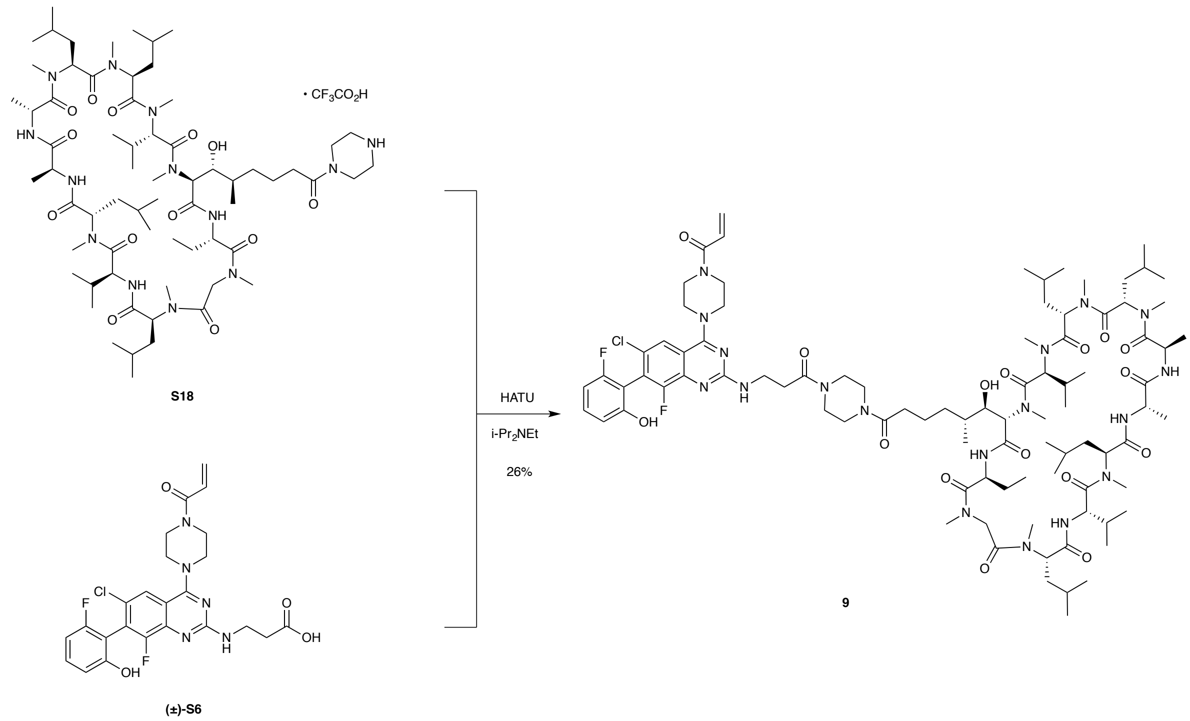


An oven dried one-dram vial was charged with **S18** (10 mg, 0.071 mmol), **(±)-S6** (4.0 mg, 0.078 mmol), DMF (0.10 mL), and a magnetic stir bar. *N,N*-Diisopropylethylamine (3.7 µL, 0.023 mmol) and HATU (3.0 mg, 0.011 mmol) were added sequentially to the reaction mixture at 23 ºC. In 1 h, LC-MS analysis showed full consumption of the starting material. The reaction mixture was diluted with 50% acetonitrile–water to a volume of 5.0 mL, and the solution was filtered through a 0.45 µM PTFE syringe filter. The filtrate was purified by reverse-phase HPLC (Waters XBridge C18 column 5 µm particle size 30 x 250 mm, 5–95% acetonitrile–water + 0.1% formic acid, 40 min, 20 mL/min) to afford the product (**9**) as a white powder (3.3 mg, 26%).

HRMS (ESI): Calcd for $(C_{90}H_{139}ClF_2N_{18}O_{16} + 2H)^{2+}$: 901.5214, Found: 901.5244.

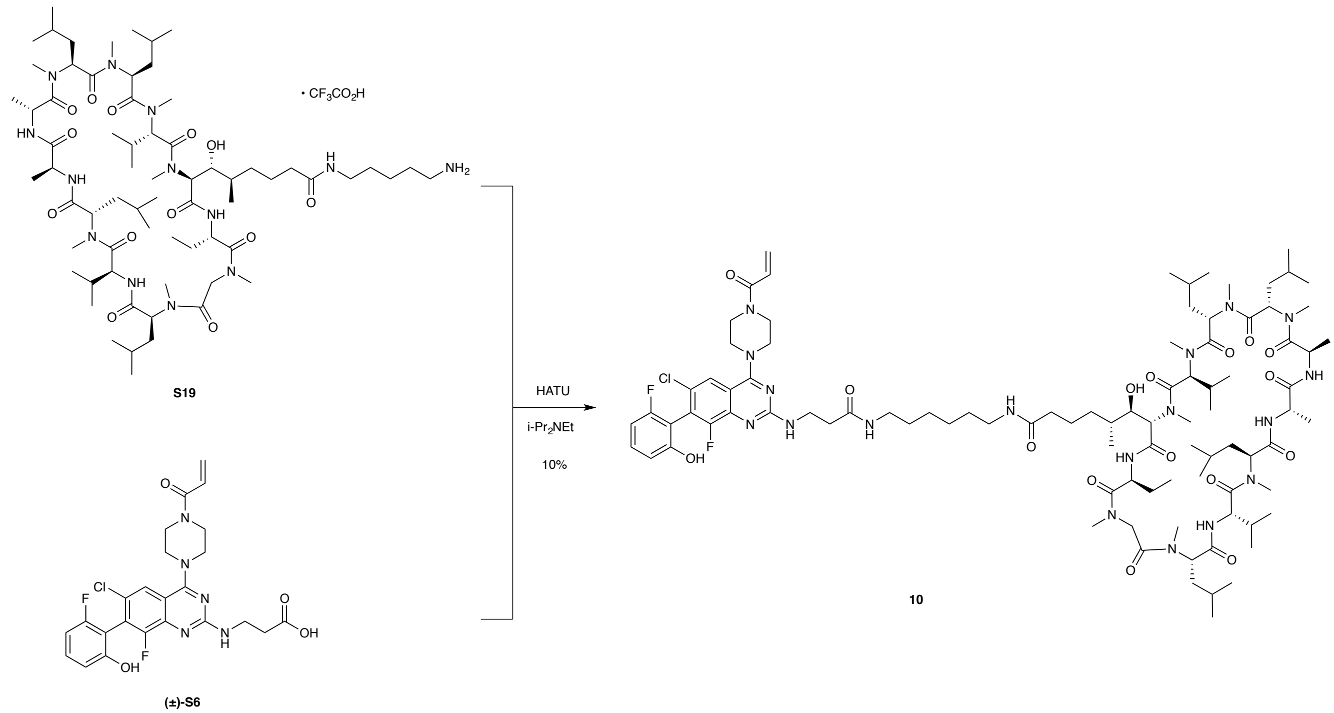

An oven dried one-dram vial was charged with **S19** (10 mg, 0.071 mmol), **(±)-S6** (4.0 mg, 0.078 mmol), DMF (0.10 mL), and a magnetic stir bar. *N,N*-Diisopropylethylamine (3.7 µL, 0.023 mmol) and HATU (3.0 mg, 0.011 mmol) were added sequentially to the reaction mixture at 23 ºC. In 1 h, LC-MS analysis showed full consumption of the starting material. The reaction mixture was diluted with 50% acetonitrile–water to a volume of 5.0 mL, and the solution was filtered through a 0.45 µM PTFE syringe filter. The filtrate was purified by reverse-phase HPLC (Waters XBridge C18 column 5 µm particle size 30 x 250 mm, 5–95% acetonitrile–water + 0.1% formic acid, 40 min, 20 mL/min) to afford the product (**10**) as a white powder (1.3 mg, 10%).

HRMS (ESI): Calcd for $(C_{92}H_{145}ClF_2N_{18}O_{16} + 2H)^{2+}$: 916.5450, Found: 916.5455.

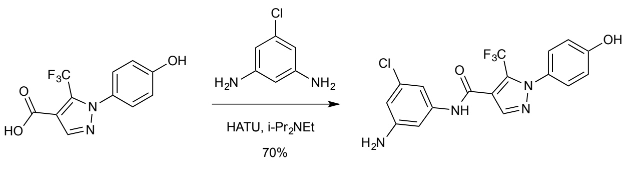

An oven-dried 20-mL vial was charged with 1-(4-hydroxyphenyl)-5-(trifluoromethyl)pyrazole-4-carboxylic acid (50 mg, 0.18 mmol), 5-Chloro-1,3-benzenediamine (131 mg, 0.92 mmol) and a magnetic stir bar. DMF (0.37mL) was added and the mixture was stirred until all reactants had dissolved. *N,N*-Diisopropylethylamine (96 µL, 0.55 mmol) and HATU (91 mg, 0.24 mmol) were added sequentially. Stirring was continued and the reaction progress was monitored by LC-MS. In 8 h, LC-MS analysis showed full consumption of the starting material and formation of one major product and one minor product. The minor product had an m/z that matched a dimer (bis-acylation). The reaction mixture was diluted with 50% acetonitrile–water to a volume of 10.0 mL, and the solution was filtered through a 0.45 µM PTFE syringe filter. The filtrate was purified by reverse-phase HPLC (Waters XBridge

C18 column 5 µm particle size 30 x 250 mm, 5–95% acetonitrile–water + 0.1% formic acid, 40 min, 20 mL/min) to afford the product (**S20**) as a white solid (51 mg, 70%).

$^1$H NMR (400 MHz, Methanol-*d*$_4$) δ 8.05 (s, 1H), 7.30 (d, *J* = 8.8 Hz, 2H), 7.03 – 6.98 (m, 2H), 6.98 – 6.89 (m, 2H), 6.52 (t, *J* = 1.9 Hz, 1H).

HRMS (ESI): Calcd for $(C_{17}H_{12}ClF_3N_4O_2 + H)^+$: 397.0679, Found: 397.0689.

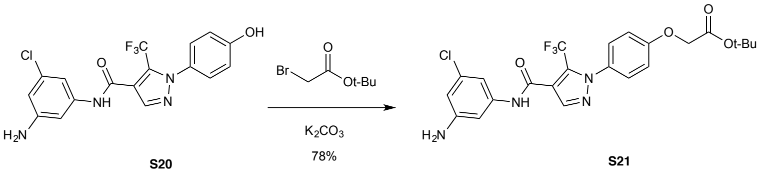


An oven-dried 1-dram vial was charged with **S20** (30 mg, 0.076 mmol), Potassium carbonate (21 mg, 0.15 mmol), DMF (0.13 mL), and a magnetic stir bar. Tert-butyl bromoacetate (11 µL, 0.076 mmol) was added via pipette, and the resulting mixture was stirred at 23 ºC. In 16 h, LC-MS indicated that the starting material had been fully consumed and two products had formed. One had the desired m/z and the other seemed to be a bis-alkylation product. The reaction mixture was partitioned between saturated aqueous sodium bicarbonate solution (5 mL) and ethyl acetate (5 mL). The layers were separated, and the aqueous layer was extracted with ethyl acetate (2 x 5 mL). The combined organic layers were dried over sodium sulfate. The dried solution was filtered, and the filtrate was concentrated. The residue was purified by column chromatography (20–80% ethyl acetate–hexanes, 4-g RediSep(R) Rf column, Teledyne ISCO, Lincoln, NE) to afford the product (**S21**) as a yellow powder (30 mg, 78%).

$^1$H NMR (400 MHz, Chloroform-*d*) δ 7.96 (s, 1H), 7.51 (s, 1H), 7.40 – 7.32 (m, 2H), 7.10 (d, *J* = 2.3 Hz, 1H), 7.04 – 6.94 (m, 2H), 6.79 (t, *J* = 1.8 Hz, 1H), 6.47 (t, *J* = 1.9 Hz, 1H), 4.58 (s, 2H), 3.83 (s, 2H), 1.49 (s, 9H).

HRMS (ESI): Calcd for $(C_{23}H_{22}ClF_3N_4O_4 + H)^+$: 511.1360, Found: 511.1376.

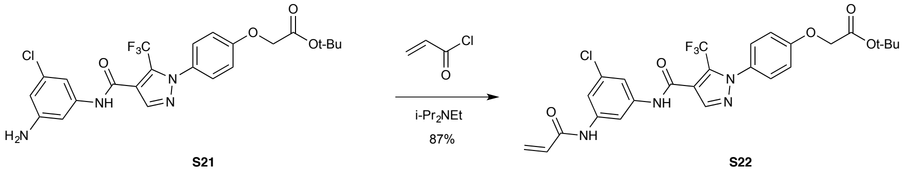


A solution of **S21** (30 mg, 0.060 mmol) in dichloromethane (0.39 mL) was cooled to 0 ºC. Triethylamine (16.37 µL, 0.12 mmol) and Acryloyl chloride (5.7 µL, 0.071 mmol) were added sequentially via syringe. The resulting solution was stirred at 0 ºC for 30 min, at which point TLC analysis (50% ethyl acetate–hexanes) indicated full conversion of the starting material to a slightly less polar spot. The reaction mixture was partitioned between saturated aqueous sodium bicarbonate solution (5 mL) and dichloromethane (5 mL). The layers were separated, and the aqueous layer was extracted with dichloromethane (2 x 5 mL). The combined organic layers were dried over sodium sulfate. The dried solution was filtered, and the filtrate was concentrated. The residue was purified by column chromatography (20–80% ethyl acetate–hexanes, 4-g RediSep(R) Rf column, Teledyne ISCO, Lincoln, NE) to afford the product (**S22**) as a yellow powder (29 mg, 87%).

$^{1}$H NMR (400 MHz, Chloroform-*d*) δ 8.81 (s, 1H), 8.33 (s, 1H), 7.94 (s, 1H), 7.82 – 7.75 (m, 1H), 7.38 (dt, *J* = 11.3, 1.8 Hz, 2H), 7.27 (d, *J* = 7.1 Hz, 1H), 7.00 – 6.88 (m, 2H), 6.32 (dd, *J* = 16.9, 1.4 Hz, 1H), 6.18 (dd, *J* = 16.9, 10.2 Hz, 1H), 5.68 (dd, *J* = 10.1, 1.4 Hz, 1H), 4.55 (s, 2H), 1.48 (s, 9H).

HRMS (ESI): Calcd for $(C_{26}H_{24}ClF_3N_4O_5 + H)^+$: 565.1466, Found: 565.1466.

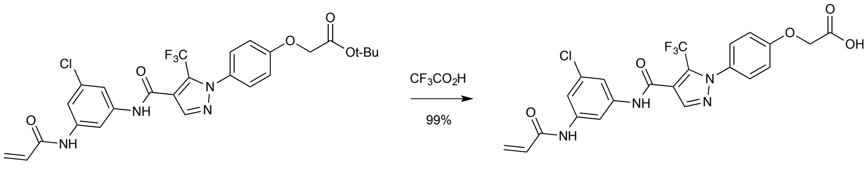


S22 S23

**S22** (29 mg, 0.050 mmol) was dissolved in 1:1 dichloromethane (0.2000mL): trifluoroacetic acid (0.2000mL) and the resulting solution was allowed to stand at 23 ºC for 1 h. The solution was then concentrated to afford the product (**S23**) as a white solid (26 mg, 99%).

HRMS (ESI): Calcd for $(C_{22}H_{16}ClF_3N_4O_5 + H)^+$: 509.0840, Found: 509.0847.

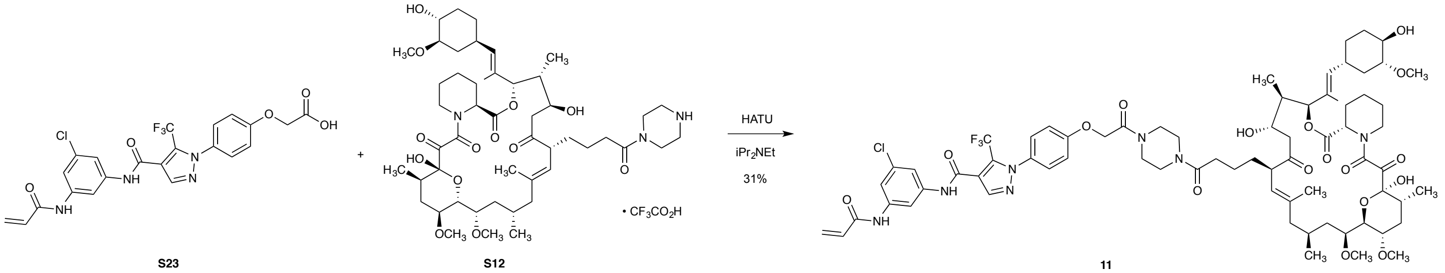


S23 S12 11

An oven-dried 1-dram vial was charged with **S23** (5.4 mg, 0.011 mmol) and **S12** (10 mg, 0.010 mmol), DMF (0.20mL) and magnetic stir bar. *N,N*-Diisopropylethylamine (8.4 µL, 0.048 mmol) and HATU (4.4 mg, 0.012 mmol) were added sequentially to the solution, and the mixture was stirred at 23 ºC while the reaction progress was monitored by LC-MS. In 1 h, LC-MS analysis showed full consumption of the FK506 starting material. The residue was diluted with 50% acetonitrile–water to a volume of 4.4 mL, and the solution was filtered through a 0.45 µM PTFE syringe filter. The filtrate was purified by reverse-phase HPLC (Waters XBridge C18 column 5 µm particle size 30 x 250 mm, 5–95% acetonitrile–water + 0.1% formic acid, 40 min, 20 mL/min) to afford the product (**11**) as a yellow solid (4.3 mg, 31%).

$^{1}$H NMR (400 MHz, Chloroform-*d*) δ 7.99 (s, 1H), 7.94 – 7.80 (m, 1H), 7.61 – 7.47 (m, 2H), 7.43 – 7.34 (m, 2H), 7.12 – 7.04 (m, 2H), 6.45 (d, *J* = 16.9 Hz, 1H), 6.29 – 6.18 (m, 1H), 5.81 (d, *J* = 10.4 Hz, 1H), 5.35 (s, 1H), 5.15 – 4.94 (m, 2H), 4.84 – 4.77 (m, 3H), 4.45 – 4.31 (m, 1H), 3.99 – 3.77 (m, 2H), 3.72 – 3.53 (m, 10H), 3.41 (s, 3H), 3.37 (s, 3H), 3.35 – 3.31 (m, 3H), 3.30 (s, 3H), 3.05 – 2.97 (m, 2H), 2.84 – 2.63 (m, 1H), 2.42 – 2.20 (m, 5H), 2.01 (d, 5H), 1.67 (s, 6H), 1.61 – 1.52 (m, 6H), 1.51 – 1.26 (m, 8H), 1.10 – 1.01 (m, 2H), 1.00 (d, *J* = 6.3 Hz, 3H), 0.93 (d, *J* = 6.8 Hz, 3H), 0.85 (d, *J* = 7.1 Hz, 3H).

HRMS (ESI): Calcd for $(C_{71}H_{93}ClF_3N_7O_{17} + H)^+$: 1408.6347, Found: 1408.6415

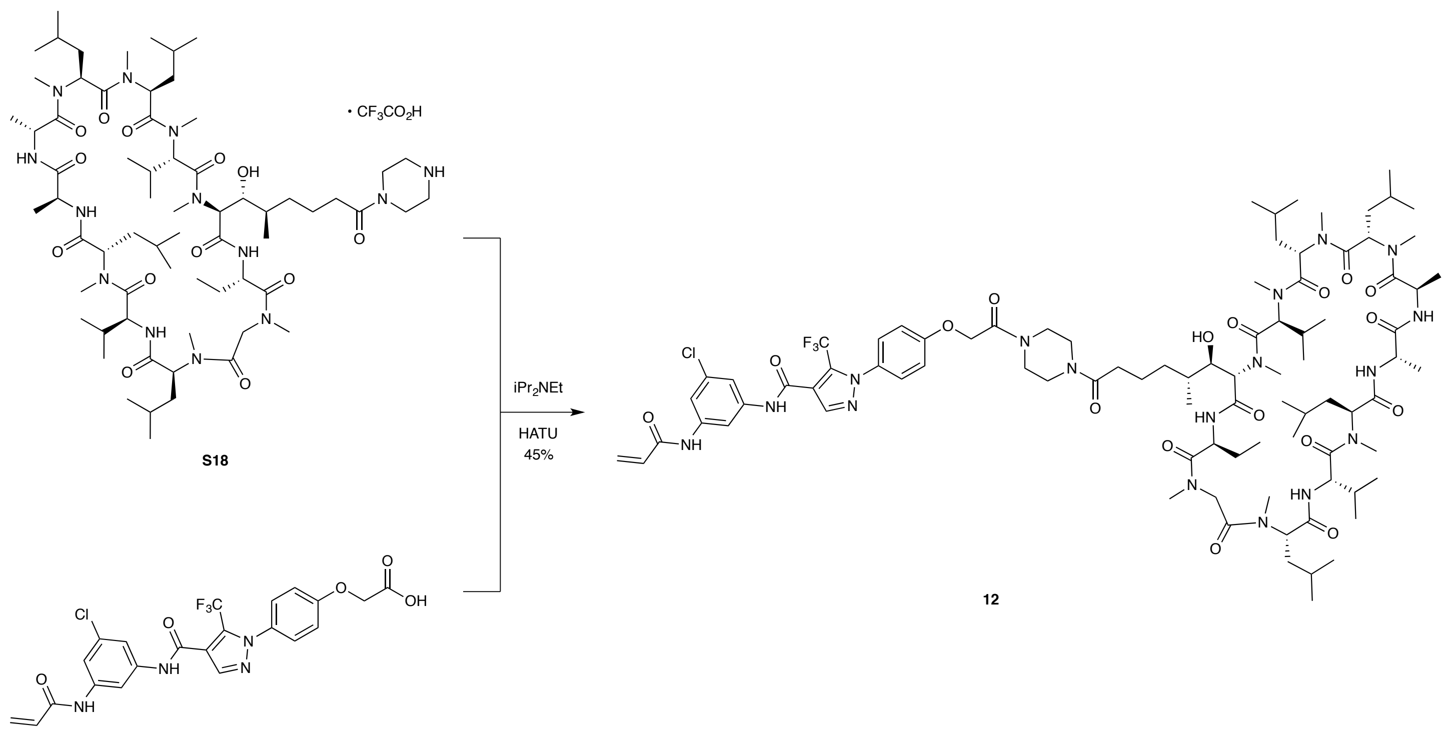


A mixture of **S18** (12 mg, 0.0085 mmol) and **S23** (4.7 mg, 0.0093 mmol) was dried by azeotropic evaporation of their suspension in benzene (1 mL). The residue was dissolved in DMF (0.20 mL). *N,N*-Diisopropylethylamine (8.4 µL, 0.048 mmol) and HATU (3.5 mg, 0.0093 mmol) were added sequentially to the solution, and the mixture was stirred at 23 ºC while the reaction progress was monitored by LC-MS. In 1 h, LC-MS showed full consumption of the 07-059 starting material. The residue was diluted with 50% acetonitrile–water to a volume of 4.0 mL, and the solution was filtered through a 0.45 µM PTFE syringe filter. The filtrate was purified by reverse-phase HPLC (Waters XBridge C18 column 5 µm particle size 30 x 250 mm, 5–95% acetonitrile–water + 0.1% formic acid, 40 min, 20 mL/min) to afford the product as a white solid (6.9 mg, 45%).

HRMS (ESI): Calcd for $(C_{88}H_{133}ClF_3N_{17}O_{17} + 2H)^{2+}$: 896.9931, Found: 896.9883.
